# Resolving Allopolyploid Origins Within the Genus *Clarkia* Using a Novel Read-Mapping and Modeling Approach

**DOI:** 10.64898/2026.09.13.751269

**Authors:** Kimmy Stanton, Mark Rausher

## Abstract

- Whole genome duplications are a common occurrence in plants, but this creates challenges for reconstructing the evolutionary history between species, especially when polyploidy is a result of hybridization. While multiple methods have been developed to try to tackle these issues, most are computationally intensive, restrictive on the number of taxa that can be evaluated, and benefit immensely from *a priori* hypotheses about the allopolyploid progenitors, rendering these methods unfeasible for many understudied polyploids.
- We present a rapid, low-cost, and computationally light method for determining the relative time of hybridization as well as the most likely progenitor species of a given allopolyploid species, including progenitors that are extinct, ancestral, or unknown.
- The method utilizes a combined approach of first mapping sequencing reads from the polyploid against a diploid pantranscriptome to generate hypotheses about possible progenitor pairs and then modeling various hybridization scenarios to estimate the likelihood of each hypothesis.
- We demonstrate the utility of our methods by identifying likely progenitors and times of origin for six allotetraploid species from the genus *Clarkia*. While the methods outlined here do not conclusively confirm the origins of these allopolyploids, they provide well-supported working hypotheses for further intensive exploration.

## Introduction

Approximately 35% of plant species are polyploids, and for angiosperms and “lower” plants (ferns, Lycophytes), respectively, approximately 15% and 31% of speciation events involve polyploidization (Wood et al. 2009). Understanding plant evolution thus necessitates characterizing the processes that generate polyploids. One challenge in doing so involves identifying the progenitors of polyploid species, specifically in the case of allopolyploid species where polyploidization is the result of hybridization between potentially distantly related taxa. For many crop and model species, progenitors have been identified only through extensive cytological, genetic and genomic analyses (e.g. Rana et a. 2004; Hurka et al. 2012; Zohary et al. 1969; Zou et al. 2015), although often uncertainties remain. These investigations have been facilitated by small numbers of potential progenitors, allowing each to be extensively examined.

Where extensive analysis is not feasible, one fully computational approach that has been developed is to reconstruct reticulating species phylogenetic networks that include allopolyploids to investigate their hybrid origins (e.g. SNaQ (Solis-Lemus and Ane 2016); Phylonet (Yan et al. 2022); Polyphest (Yan et al. 2024)). However, these approaches have their own limitations (Yan et al. 2024). First, these methods typically require a set of orthologous genes to be identified and that the subgenomes be phased before the analyses can be applied. In addition, *a priori* assumptions must often be made about the progenitors of subgenomes, which is often not ideal for systems where such information is not readily available. Finally, these methods are generally computationally intensive and scale in a way that is problematic for large numbers of loci or taxa (Blischak et al. 2020).

For evolutionary biologists, this type of extensive analysis of allopolyploid progenitors is often prohibitively expensive, both financially and effort-wise, especially if a large number of possible progenitors need to be considered. Consequently, it would be helpful to have a method that is relatively inexpensive and that requires relatively little effort to identify likely progenitors that can be further analyzed intensively. An alternative approach that has been suggested is to use read mapping of polyploid sequences to diploid genomes to determine which candidate progenitor provide the best matches to a given read (Wang et al. 2021). Such an approach is computationally simple, rapid, does not require any *a priori* hypotheses, and can provide a well-supported hypotheses regarding the progenitor species for a given allopolyploid, which is necessary for more intensive downstream genomic, phylogenetic, and cytological analyses. This read-mapping method can be especially useful in cases of recent hybridization between extant species, in which the subgenomes have had little time to diverge from the progenitor sequences. However, due to the stochasticity of substitutions, the influence of incomplete lineage sorting, and the age of the polyploid (with older polyploidization events presenting further challenges for progenitor discovery), read alignments (hereafter termed “matches” or “best hits”) can also be distributed among many potential diploid species, making interpretation complicated.

One solution to this problem is to model substitutions in both diploid and polyploid subgenomes across a highly supported species phylogeny to estimate the expected proportions of best hits of polyploid reads to each possible diploid progenitor. If one constructs models for different pairs of likely progenitors, based on hypotheses generated from the read mapping data, and additionally allows hybridization times to vary, the model producing the closest match to the observed distribution of hits corresponds to the most likely progenitors. Two major benefits to this combined read mapping and modeling approach are the ability to determine an approximate time of hybridization (either absolute or relative) and to assess whether one of the progenitors is currently unknown, ancestral, or extinct.

Here we describe an implementation of the combined read-mapping and modeling approach and use it to identify the most likely diploid progenitors of six allotetraploid species in the angiosperm genus *Clarkia*, as well as to determine the relative timings of hybridization. We show that the read mapping approach is a powerful tool for filtering down to a few taxa/lineages that are most likely involved in the hybridization event. We also show that evolutionary modeling focused on potential hybridizing pairs from among these few taxa can generate well-supported hypotheses about the diploid progenitors of allotetraploids, regardless of whether the progenitor is unknown.

## Methods

### Plant Species, Transcriptome Data, and Diploid Phylogeny

As reported previously in Stanton et al. 2026, petal transcriptomes were obtained from 28 of the 32 diploid species in the genus *Clarkia* and from most subspecies of the 10 polyploid species in the genus. Seed sources, sample names, and herbarium vouchers (if available) for the 38 plant lines used in this study are listed in Supplementary Table SIV.1. Four diploid species were not sampled due to difficulty obtaining seeds: *C. springvillensis, C. modesta, C. jolonensis*, and *C. heterandra*. Here we report results for six of seven tetraploid species: *C. rhomboidea, C. pulchella, C. similis, C. delicata, C. tenella*, and *C. davyi*. One tetraploid, *C. gracilis*, was excluded because of the polytomy between the three likely progenitor species (Supplementary Fig. S1.1D,E; Sianta and Kay 2022). The three hexaploid *Clarkia* species (*C. affinis, C. prostrata*, and *C. purpurea*) were not included in this analysis because our current modeling method is not designed to accommodate hexaploids.

RNA was prepared and sequenced, and transcriptomes were assembled for all samples as outlined in Stanton et al. (2026). In brief, RNA was extracted from petal tissue collected one day before flowering, and libraries were prepared using the Kapa Stranded RNA-seq Kit (Roche) and barcoded with NEBNext Unique Dual Index Primer Pairs (NEB, Cat#E6442). Libraries were sequenced on an Illumina HiSeq 4000, Illumina NovaSeq 6000, or Illumina NovaSeq X Plus platform at the Duke University Sequencing Core. Reads from each sample were trimmed with TrimGalore v.2.2.0 (https://github.com/FelixKrueger/TrimGalore) and *de novo* assembled into a transcriptome using Trinity v.2.15.2 software (Grabherr 2011).

Also as previously reported in Stanton et al. (2026), phylogenetic reconstruction of the *Clarkia* diploid species tree was performed using two different methods (using ASTRAL-III (Zhang et al., 2018) and by maximum likelihood using RAxML-ng (Kozlov et al. 2019)), yielding trees with excellent support at the species level. Both approaches produced the same tree topology, with two minor differences: the placement of *C. dudleyana*, and the relationships among *C. rubicunda, C. franciscana*, and *C. amoena*. For this study, we used an ultrametric tree based on the RAxML tree (Kozlov et al. 2019) (Supplementary Fig. SI.1).

### Polyploid Read Mapping

In order to identify the putative diploid parents of the six tetraploid *Clarkia* species, a pantranscriptome was constructed by concatenating the transcriptomes from the 28 diploid species examined in this study, using only one representative sample per species and excluding subspecies. The trimmed sequencing reads from each polyploid were then aligned to this pantranscriptome using salmon v.2.1.1 (Patro et al. 2017). Any gene in the concatenated pantranscriptome set that received a number of alignments greater than 10 TPM was scored as 1 “match” or “best hit” for that gene for the corresponding diploid species. These best hits were summed for each species to produce a vector of number of best hits, *n* = {*n*_*1*_, *n*_*2*_, …, *n*_*28*_}, where *n*_*i*_ is the total number of best hits to diploid species i. The proportions of best hits (PBH) for each species was then calculated as follows and provided in (Supplemental Table SIV.2):

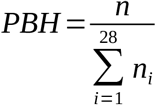

We term this vector the “observed PBH spectrum” and present it as a histogram (e.g. Fig. 1). We use this index instead of the proportions of reads aligning to each diploid species to minimize effects of different genes being expressed at different levels. Estimates of observed PBH values for replicate plant lines within a species were extremely similar, even if the replicate lines came from morphologically divergent subspecies, so PBH values were averaged across replicates, when available, for all subsequent analyses.

**Figure 1.**
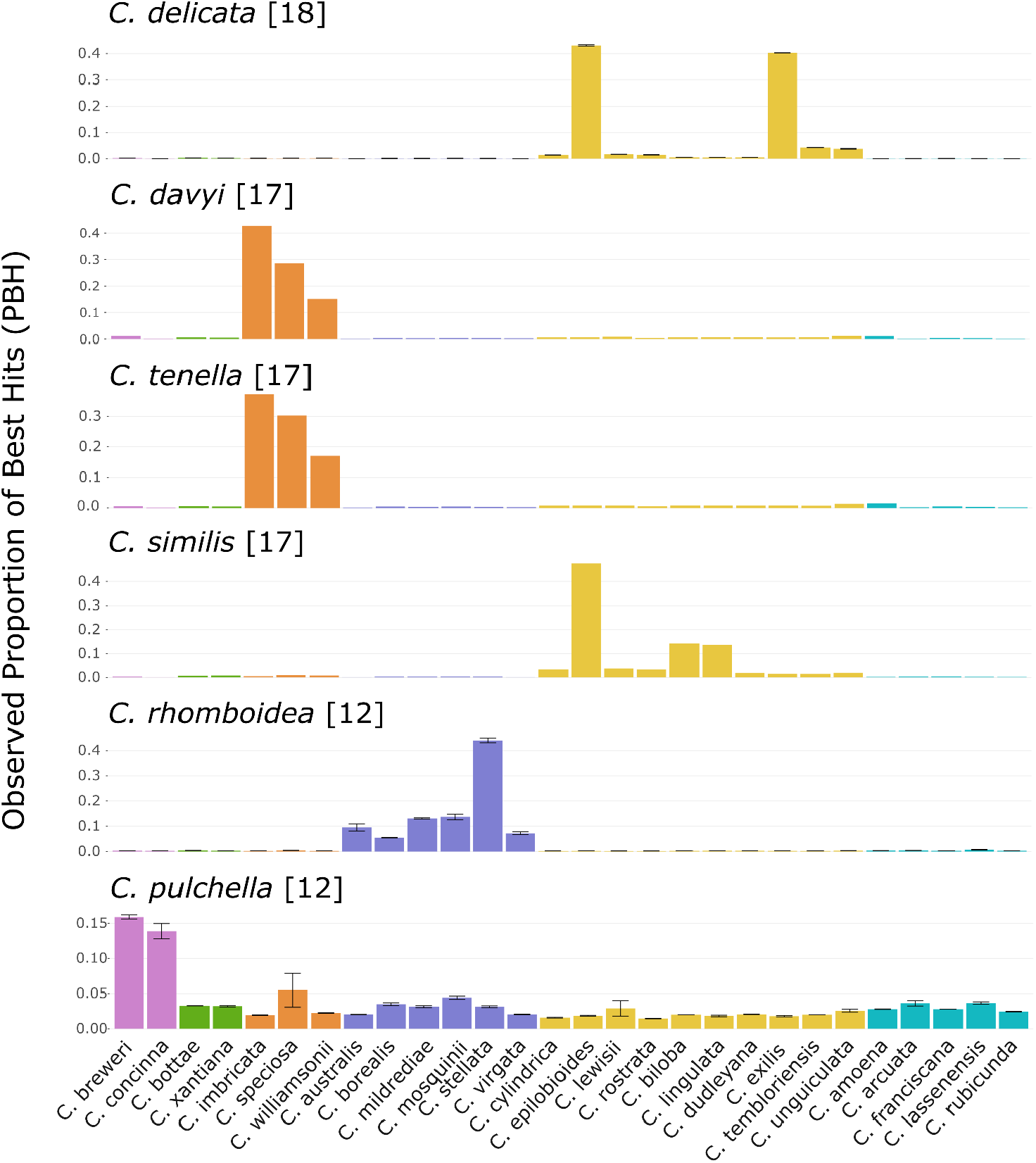
Observed proportion best hits (PBH) spectra for reads from the seven *Clarkia* tetraploid species (labeled above each graph) to the diploid pantranscriptome. Chromosome number for each tetraploid is noted next to the species name in brackets. Standard error bars are included for the three species with replicate individuals: *C. delicata, C. pulchella*. and *C. rhomboidea*. Colors represent *Clarkia* Sections as described by Wagner et al. 2007: Pink = Section *Eucharidium*, Green = Section *Fibula*, Orange = Section *Godetia*, Purple = Section *Myxocarpa*, Yellow = Section *Phaeostoma*, Blue = Section *Rhodanthos*.

### Modeling

While interpreting the PBH spectra seems relatively straightforward if most hits are allocated to just two species (for tetraploids), many tetraploids we examined have relatively high proportions of hits to more than 2 species. Consequently, in order to infer the actual progenitors, we performed evolutionary simulations. These simulations are described in detail in Supplementary Material II. Here we provide an overview.

Our modeling approach requires an ultrametric tree. As described in the supplementary material (Supplementary Material I), we used the chronos program within the package ape for R (Paradis et al 2004) to convert the RAxML tree to an ultrametric tree (Figs. SI.1, SI.2). Because this conversion requires assumptions about the tempo of substitutions, we constructed trees under a variety of different assumptions (e.g. strict molecular clock, relaxed clock, correlated clock, discrete clock) and found no reason to reject the assumption of a strict clock (Supplementary Material I).

Genomes were simulated as 40 genes of 750 bp each (30,000 bp total). Genomes were represented by a vector of numbers (0, 1, 2, or 3) representing the four individual DNA nucleotides. The genotype at the root of the tree was initialized as a vector of all 0’s. The number of substitutions along each branch, designated generically as “node A → node B”, was simulated as a Poisson process with two parameters: (1) the length of the branch, in relative time units (“RTU”; time from root to tip is assumed to be 1 RTU), and (2) the rate parameter λ = 0.1913893 estimated from the ultrametric tree (Supplementary Material I). Starting with the genome at node A, the substitutions were randomly allocated to individual nucleotide sites and the change at each site (e.g., 0→1, 4→2, 3→0) was determined randomly. The new genome was stored as the genome associated with Node B. This produced a set of simulated genomes for each node in the tree, including the tip nodes (diploid species).

Each simulation also hypothesized a hybridization between two branches at a specific relative time before present, *T*_*H*_. These branches could correspond to individual species, the common ancestor of two or more species, or, in some cases, a postulated unknown (e.g. extinct, unidentified, or one of the four diploid *Clarkia* species not included in our analysis). These branches, which we refer to generically as “node X → node Y”, were broken into two segments: (1) node A → node H, and (2) node H → node B. From node H, evolution of one subgenome of the hybrid was simulated in the same manner as evolution along the tree branches, with the length of the branch being equal to the time of hybridization in RTU. These calculations produced two genomes corresponding to the two subgenomes of the tetraploid species.

A detailed explanation of how we calculated the simulated proportion best hits of each diploid species is provided in Supplementary Material II. Here we provide a brief summary: we first divided the genome into 40 750bp segments corresponding to “genes”. We further subdivided each “gene” into 5 150bp-segments, corresponding to the tetraploid RNA “reads” from the transcriptome analysis. We designate each read by *r*_*ijk*_, indicating that it is a read from segment *i* (*i ∈* {1,2,3,4,5 }) of “gene” *j* ( *j ∈* {1,2, …, 40 })) in tetraploid subgenome *k* (*k ∈* {1,2 }). For each read (one per segment per gene) we calculated a set of distances between that read and the corresponding segment and gene in each diploid genome, where distance is defined as the proportion of nucleotides that differ (= Hamming Distance). Each read *r*_*ijk*_ is then assigned as an alignment to gene *k* of the diploid with the lowest difference. If there is more than one diploid with the lowest difference, the alignment is assigned randomly.

Unlike when mapping actual reads, in our model we know which “genes” are orthologous. Consequently, for each “gene” *j*, the above procedure yields a vector of alignments (“Best Hits”) of the form

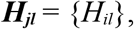

where each element *H*_*il*_ = 1 if at least one “read” from either subgenome aligns to gene *j* of diploid species *l ∈* {1,2, … 26} and 0 otherwise. Schematically, a hypothetical result could be

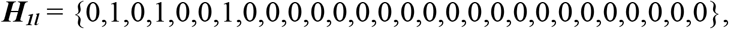

i.e. reads from gene 1 aligned to segments diploid species 2, 4 and 7.

Summing over genes produces a vector of simulated number of best hits for the tetraploid to each of the diploid species:

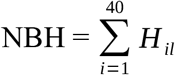

This vector is then divided by the total number of best hits to yield a vector of proportion of best hits (PBH) to each diploid species:

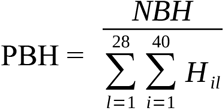

This process was repeated 1000 times to produce 1000 simulated PBH vectors (simulation replicates), which were then averaged to obtain a mean, 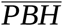. When plotted as a histogram, we term this the “simulated PBH spectrum”.

Because we used RNA sequencing data, the complication of subgenome dominance arises, where more genes tend to be lost from, or tend to be more highly expressed in, one tetraploid subgenome compared to the other (Bird et al. 2018). The procedure outlined above assumes that both subgenomes contribute equally to the simulated PBH spectrum. However, we have modified this procedure to account for different levels of subgenome dominance as described in Supplementary Material II.

We assessed how well a simulation reproduced the observed proportion hits in two ways. First, we compared the observed and simulated PBH spectra visually. This comparison allowed determination of whether a simulation was a poor fit to the data. Second, we calculated the sum of squared differences between the observed and simulated spectra, which allowed us to compare the relative fit of multiple simulations with different hybridizing species and different hybridization times. This procedure allowed us to identify the set of parameters (i.e. hybridization time, subgenome dominance, and unknown species split time) that provided the best fit for each prospective progenitor pairs. To account for error associated with simulations, we then conducted 50 replicate simulations using those parameters to obtain a frequency distribution of SS values for each best fitting spectrum. If the distributions for two progenitor pairs did not overlap, we inferred that the more likely pair was the one with the lowest mean SS. By contrast, if the distributions overlap, we concluded that neither pair was more likely than the other.

## Results

### Tetraploid read mapping

Mapping of RNA sequencing reads from the six tetraploid *Clarkia* species evaluated in this study to the diploid pantranscriptome in most cases produced a small number of candidate progenitors based on the observed proportion best hits (PBH) spectrum (Fig. 1; values given in Supplementary Table SIV.2). For most polyploids, there are two or three diploid species that account for most of the hits, with other species accounting for much lower proportions. But, as can be seen in Figure 1, for all tetraploid species except *C. delicata*, more than two diploid species receive a substantial proportion (>5%) of best hits. Because each tetraploid hybrid had only two parental species, it is *a priori* not clear how to interpret these results. Consequently, we undertook modeling analyses to infer the identity of the most likely diploid progenitors for these tetraploid species.

### Simulations for a clear, biparental case

*C. delicata* was one of the few cases where the PBH spectrum was clearly dominated by only two diploid species, *C. exilis* (40.3%) and *C. epilobioides* (43.1%), which indicates that these two species are the most likely candidates for the progenitors of *C. delicata*. We simulated the evolution of hybrids produced by 5 prospective pairs of diploid species (Fig. 2A), with most pairs evaluating *C. epilobioides* hybridizing with either *C. exilis* or a closely related species, and compared the PBH spectra of these simulated hybrids to the observed spectrum. The best-fitting model corresponds to hybridization between *C. exilis* and *C. epilobioides* at 0.19 RTU before present (Fig. 2B,C). The simulated PBH spectrum for this model is qualitatively very similar to the observed spectrum whereas the other models that were evaluated had SS values that are at least 20 times higher, as well as simulated PBH spectra that were qualitatively poor fits to the observed spectrum (Supplementary Fig. SIII.1).

**Figure 2.**
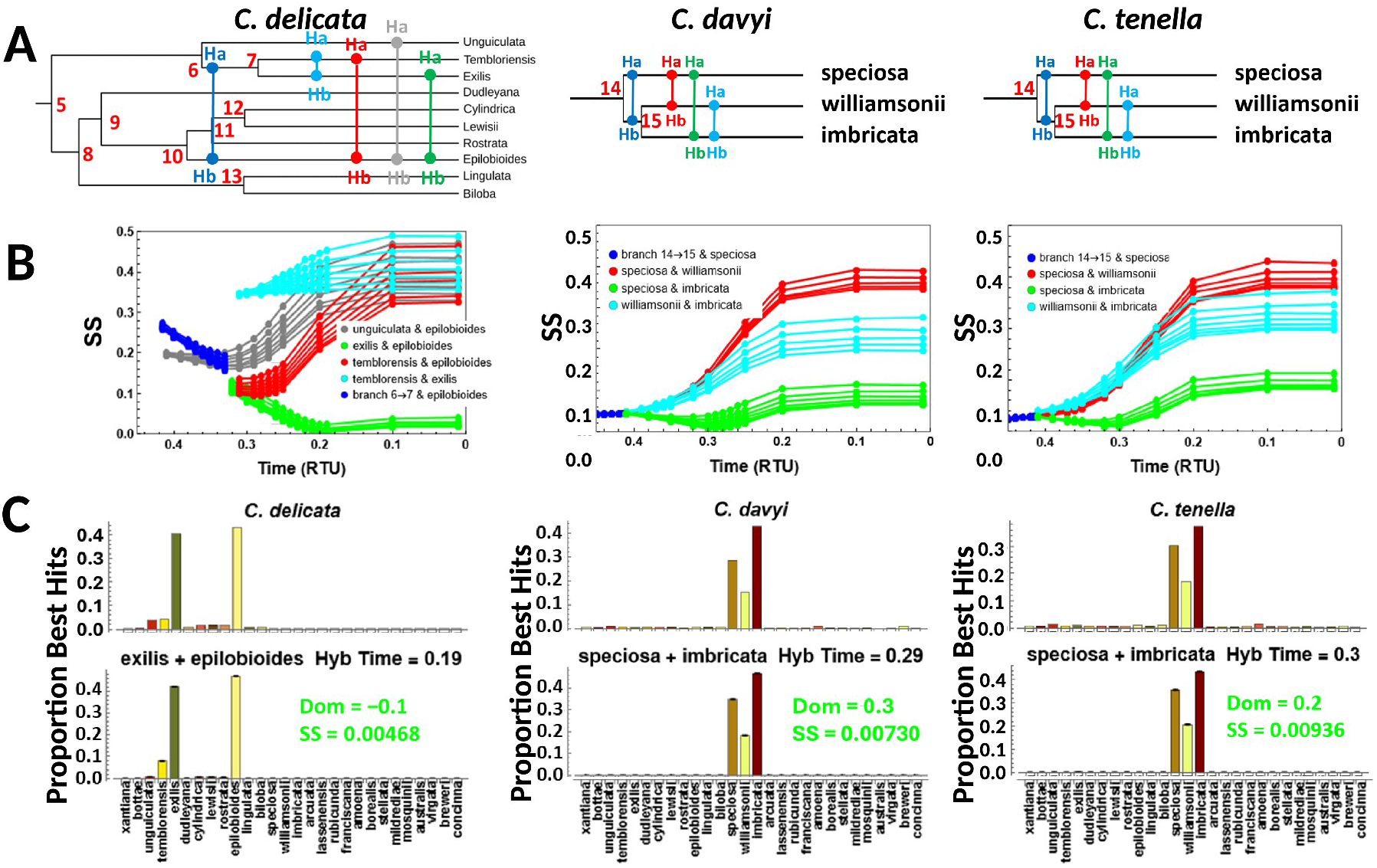
Analysis of potential diploid progenitors for the tetraploids *Clarkia delicata* (column 1), *Clarkia davyi* (column 2), and *Clarkia tenella* (column 3) via simulations. **A**. Phylogenetic relationships for the pairs of hybridizing species examined. Red letters/numbers: node labels. **B**. Sum of Squares (SS) for different pairs of hybridizing species as progenitors of each tetraploid at different possible hybridization times. Replicated lines of the same color correspond to simulations for a given hybridizing pair with different levels of subgenome dominance. Hybridizing species pairs are given in the key; “6→7”: common ancestor of *C. temblorensis* and *C. exilis; “*14→15”: common ancestor of *C. williamsonii* and *C. imbricata*. **C**. Observed PBH spectrum (top), as shown in Figure 1, compared to the simulated PBH spectrum (bottom) for the best performing model. Simulated PBH spectra for lower performing models, and for models under different subgenome dominance levels are displayed in Supplementary Figs. SIII.1-6. Spectra correspond to a combination of simulation parameters (hybridization time (“Hyb Time”) and subgenome dominance (“Dom”) level) with the lowest SS for a given progenitor species pair.

### Simulations to evaluate polyploidy from a single origin

There has long been speculation that the tetraploids *C. tenella* and *C. davyi* could possibly have a tetraploid ancestor in common due to a shared number of chromosomes and strong morphological similarities between the two species (Lewis and Lewis 1955, Raven and Lewis 1959). While we didn’t simulate this claim explicitly, we wanted to evaluate whether simulations for each tetraploid refuted this hypothesis, as well as identify possible progenitor species for these two tetraploids.

Both *C. davyi* and *C. tenella* had the highest observed PBH to the same three species, *C. imbricata, C. speciosa*, and *C. williamsonii*, the only three diploid members of the *Godetia* Section (Wagner et al. 2007), indicating that a hybridization event involving two members of this clade is the most likely origin for these tetraploids (Fig. 1). The PBH values also showed a very similar pattern between the two species, while PBH for all other diploid species outside of the *Godetia* Section was below 2.0%, making it unlikely that a species outside of this Section was involved.

To model possible hybridization events that produced *C. davyi* and *C. tenella*, we simulated the evolution of hybrids produced by pairwise crosses between *C. imbricata, C. williamsonii*, and *C. speciosa* (Fig. 2A). Among the four prospective progenitor pairs, a hybridization event between *C. speciosa* and *C. imbricata* produced the best-fitting simulated PBH spectrum and yielded the lowest value of the Sum-of-Squares (SS) difference when compared to the observed PBH spectrum for both *C. davyi* and *C. tenella* (Fig. 2B,C; Supplementary Figs. SIII.3-6). The best-fitting model indicates that hybridization occurred between *C. speciosa* and *C. imbricata* at approximately 0.29 relative time units (RTUs) before present for *C. davyi* and 0.30 RTUs before present for *C. tenella*. This difference is almost certainly not significant, and thus we fail to reject the hypothesis that the two tetraploid species may be derived from a single hybridization event.

### Simulations involving unknown taxa

The observed PBH spectra for both *C. similis* and *C. rhomboidea* showed one diploid receiving a majority of the PBH, while a small handful of related diploids received between 5-15% of the PBH (Fig. 1). This pattern led us to explore through simulations the possibility that an unknown taxon (either extinct or unsampled) was involved in the origins of these two tetraploids.

In the case of *C. similis, C. epilobioides* received a PBH of 47.5%, making it a clear candidate as one of the potential progenitors, while only two other diploids received a PBH higher than 4%, *C. lingulata* (13.7%) and *C. biloba* (14.2%). Consequently, we simulated hybridizations between species pairs that included *C. epilobioides* hybridizing with *C. biloba, C. lingulata*, their common ancestor, or an unknown species sister to *C. biloba* and *C. lingulata* (Fig. 3A). Three of these simulated hybridizations have the best models with the lowest, essentially indistinguishable, SS values and exhibit simulated PBH spectra qualitatively similar to the observed spectrum: *C. epilobioides* x *C. lingulata, C. epilobioides* x *C. biloba*, and *C. epilobioides* x unknown species (Fig. 3B,C, Supplementary Figs. SIII.7A,B). The simulated PBH spectrum for the *C. epilobioides* x unknown species appears to be the closest match to the observed spectrum, with relatively equal hits to *C. lingulata* and *C. biloba*, while the two other hybridizations produce PBH spectra that are qualitatively different from the observed (Fig. 3C). The most likely progenitors were thus *C. epilobioides* and an unknown species.

**Figure 3.**
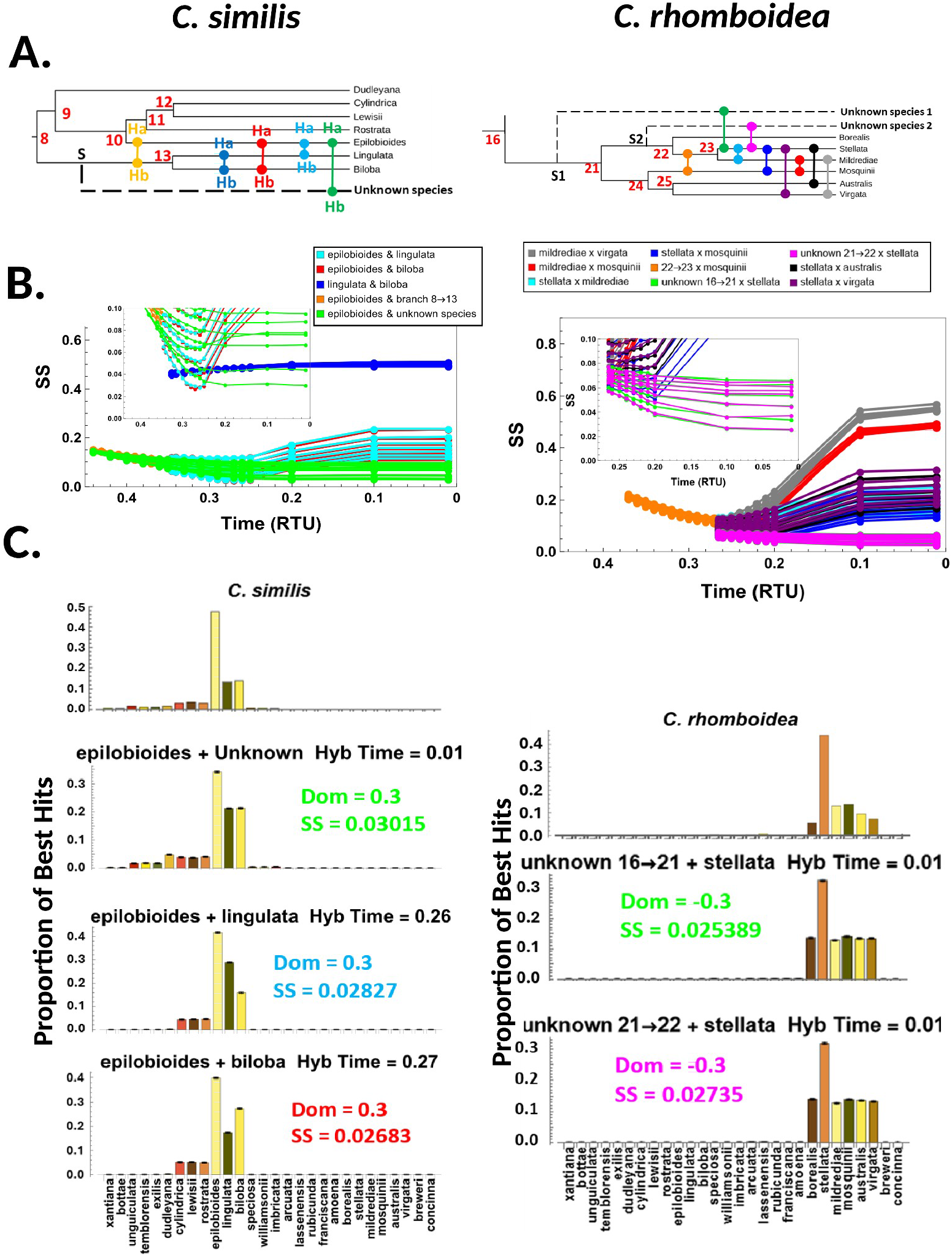
Analysis of diploid progenitors for the tetraploids *Clarkia similis* and *Clarkia rhomboidea* via simulations. Panels laid out as in Figure 2: **A**. Phylogenetic relationship for the pairs of hybridizing species examined. **B**. Sum of Squares (SS) for different pairs of hybridizing species as progenitors at different possible hybridization times. For hybridization involving an unknown species, the lines correspond to the curve for each subgenome dominance value with the lowest SS (i.e. correspond to lines in Supplementary Figs. SIII.9,12,13). **Inset**. Enlargement of panel B for times 0.4 – 0 RTU for *C. similis* and for times 0.3 – 0 RTU for *C. rhomboidea*. **C**. Observed PBH spectrum and simulated PBH spectra for different hybridizing species pairs. Simulated PBH spectra for lower performing models, and for models under different subgenome dominance levels are displayed in Supplementary Figs. SIII.7-13.

In the case of *C. rhomboidea, C. stellata* received 44% PBH, making it a clear candidate as one of the potential progenitors, while the remaining five diploid species from Section *Myxocarpa* had PBH that ranged between 5.5% and 13.7%. To model possible scenarios, we focused on pairwise simulations between extant diploid species within the Section, involving the most recent common ancestor of certain taxa, and involving two unknown species that diverged from the clade at different times (Fig. 3A). Simulated hybridizations between *C. stellata* and each of the two unknown species exhibited the smallest and indistinguishable SS values as well as qualitatively similar spectra to the observed (Fig. 3B,C; Supplementary Fig. SIII.10A,B). The remaining hybridizations examined all have even larger SS values and provide qualitatively worse fits in all cases, and the previously postulated hybridization between *C. mildrediae* and *C. virgata* is included among these cases (Lewis and Lewis 1955; Supplementary Fig. SIII.10C).

### Simulations for an older polyploid

Unlike the other polyploid species analyzed so far, for *C. pulchella* no single species received a majority of the PBH. *C. breweri* and *C. concinna* have the largest, and roughly equal, PBH values (15.9% and 13.9% respectively); however, together they received less than 30% of the best hits, with all other species receiving between 1.5% and 5.5% of best hits. This pattern is unique among the analyzed tetraploids, and is consistent with *C. pulchella* arising from an older hybridization event, which has been previously postulated (Sytsma 1990).

Based on this reasoning, we examined eight different possible progenitor pairs (Fig. 4A), most of which involve hybridization between the ancestor of the *Eucharidium* Section (containing the diploid species *C. brewerii* and *C. concinna* and denoted by branch R⟶26) and an early branch of the tree, or between those two species and an unknown, and probably extinct, species that branched off very early in the phylogeny. Among these potential progenitor pairs, hybridization involving the unknown species and the ancestor of the *Eucharidium* Section (R→26) has the lowest SS and produces a simulated PBH spectrum qualitatively similar to the observed spectrum (Fig. 4). However, several of the other hybridizing pairs involving the common ancestor of major clades of *Clarkia* also have SS values that are low and indistinguishable from the simulations involving the unknown species and the ancestor of the *Eucharidium* Section (R→26), though their simulated PBH spectra are not as good of a fit qualitatively.

**Figure 4.**
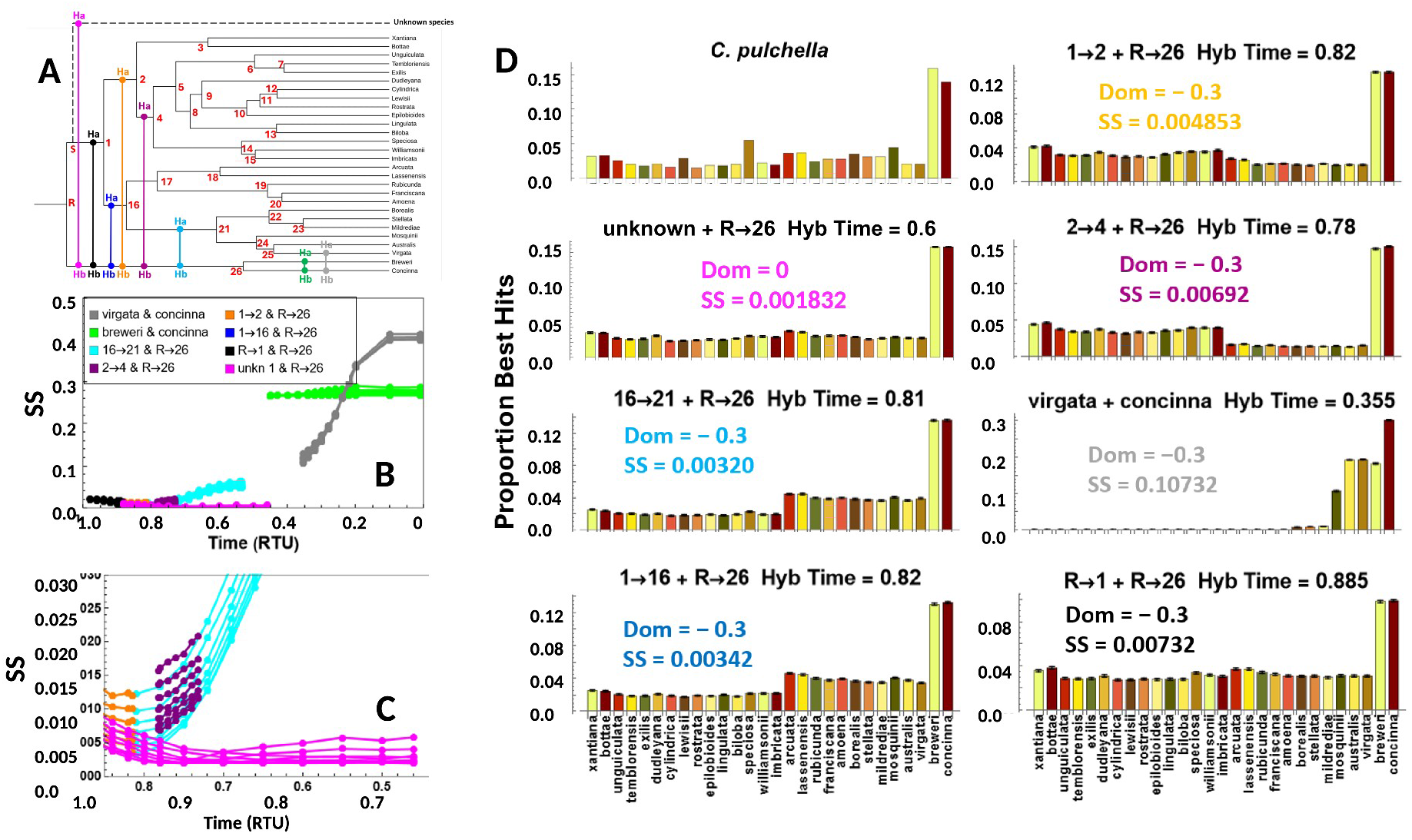
Analysis of diploid progenitors for the tetraploid *Clarkia pulchella* via simulations. **A**. Phylogenetic relationship for the pairs of hybridizing species examined. Nodes numbered in red. **B**. Sum of Squares (SS) for different pairs of hybridizing species as progenitors of *C. pulchella* at different possible hybridization times. Replicated lines of the same color correspond to simulations for a given hybridizing pair but with different levels of subgenome dominance (see Supplementary Fig. SIII.15). For hybridization involving an unknown species, the lines correspond to the curve for each subgenome dominance value with the lowest SS (i.e. correspond to lines in Supplementary Fig. SIII.16). Hybridizing species are given in key. “16→21”: common ancestor of the *Myxocarpa* Section, “2→4”: common ancestor of the clade containing Sections *Phaeostoma* and *Godetia*, “1→2”: common ancestor of the clade containing Sections *Phaeostoma, Godetia*, and *Fibula*, “1→16”: common ancestor of clade containing Sections *Rhodanthos* and *Myxocarpa*, “R→1”: common ancestor of all *Clarkia* species excepting *C. breweri* and *C. concinna*, “R→26”: common ancestor of *C. breweri* and *C. concinna*. **C**. Enlargement of panel B for times 1 – 0.65 RTU. **D**. Observed PBH spectrum (top left) and simulated PBH spectra for different hybridizing species pairs. Spectra correspond to a combination of parameters (hybridization time (“Hyb Time”) and subgenome dominance (“Dom”) level) with the lowest SS for a given species pair.

**Figure 5.**
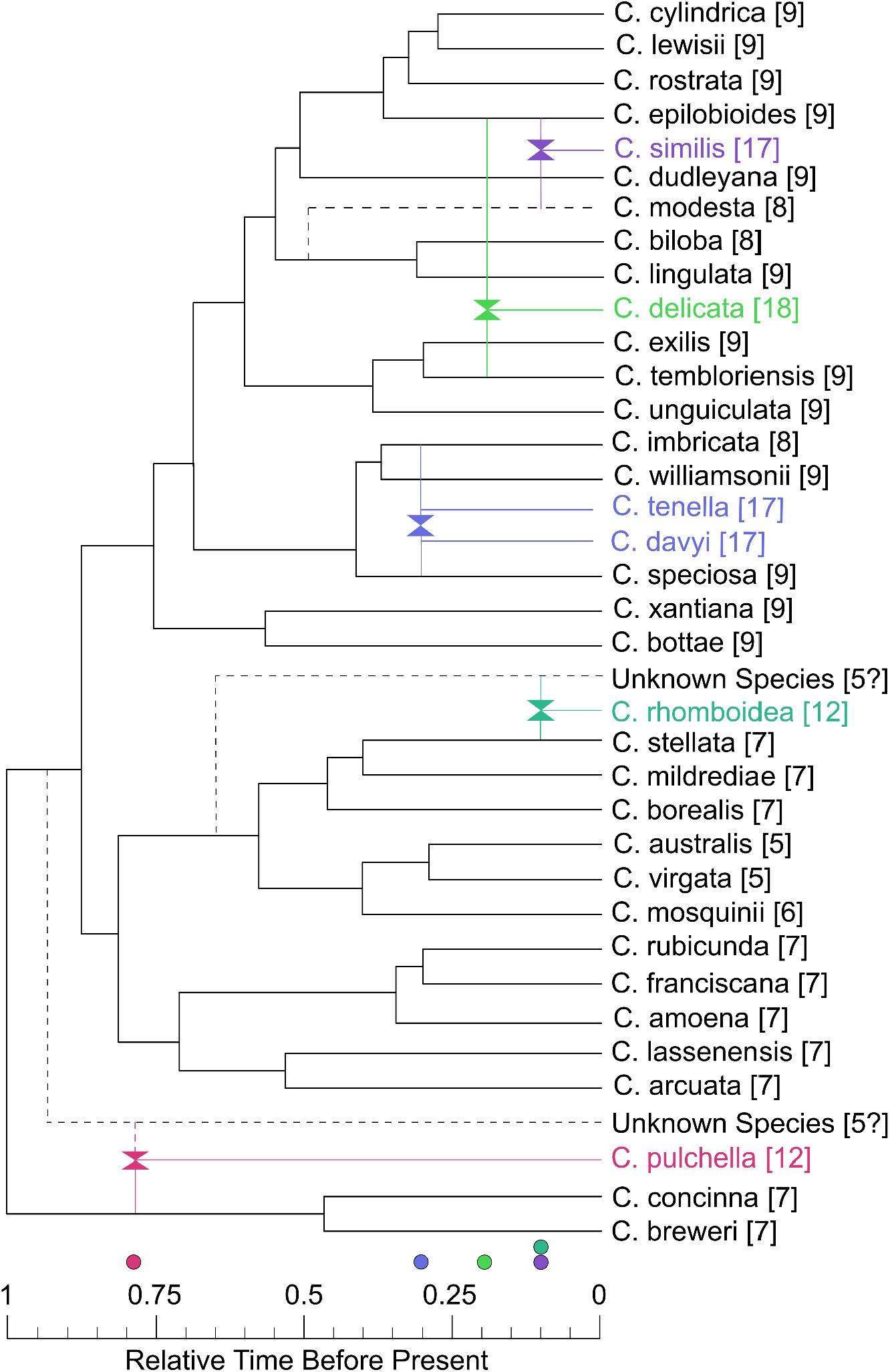
Phylogenetic network illustrating putative progenitor diploid species for the six of the allotetraploid *Clarkia* species based on the results of our read-mapping and modeling method. Dashed lines show uncertain relationships, as well as potential unknown/extinct species that were inferred from our simulations. Chromosome number for each species is reported in brackets next to the species name. Color of each species corresponds to the marker on the scale bar below marking the estimated time of hybridization, in relative time units (RTU), for the best performing models. Hybridization time is marked as 0.1 for both *C. similis* and *C. rhomboidea*, which is the middle of the likely range of hybridization (between 0.2 and 0) for both species. Diploid tree depicted was adapted from the ultrametric tree (Supplementary Fig. SI.1).

Simulated hybridizations between extant species of *Clarkia*, including *C. brewerii* x *C. concinna* and the previously postulated progenitors *C. virgata* x *C. concinna*, produce PBH spectra that are very poor qualitative matches to the observed spectrum and SS that are substantially higher than and distinct from that of other potential progenitor pairs (Fig. 4; Supplemental Fig. SIII.14). Although we can’t realistically distinguish among the best potential models, it seems highly likely that the hybridization that produced *C. pulchella* occurred early in the history of *Clarkia* and involved lineage R→26 and another early lineage (e.g. unknown species, 16→21, 1→16, 1→2, or 2→4).

## Discussion

In this investigation, we have described a combined read-mapping and modeling approach for identifying the most likely progenitors of an allotetraploid species from among a reasonably large number of possible diploid candidates. The read-mapping method we have utilized is fast, computationally light, does not require any *a priori* hypotheses, and generates hypotheses regarding progenitors that can then be further tested by more computationally intensive modeling or phylogenetic network methods. A similar read-mapping approach has been used elsewhere by Wang et al. (2021), who performed no evolutionary modeling and assumed that for tetraploids, the two extant diploid species with the greatest PBH were progenitors. Our approach represents an advance over theirs by allowing for the possibility that hybridization involved lineages corresponding to internal branches of the phylogenetic tree or lineages that are unknown (unsampled or extinct). Additionally, our modeling approach provides estimates of the approximate time of hybridization.

We have demonstrated the utility of this type of combined read-mapping and modeling approach by evaluating a wide variety of situations—including cases of recent hybridization, single hybrid origins, hybridization involving unknown taxa, and a case of relatively old hybridization—for six tetraploid species in the genus *Clarkia*.

### Implications for Clarkia

In general, our analyses are consistent with earlier evidence in that we identify potential progenitors that are closely related to, or ancestral to, previously hypothesized progenitors. However, in many cases we fail to support the exact species pairs previously suggested to be the diploid progenitors, and instead we provide updated hypotheses for the origin of six tetraploids in the genus *Clarkia* below (Figure 8).

It has long been proposed that *C. delicata* was derived from a hybridization between *C. unguiculata* and *C. epilobioides* based on morphological and cytological evidence (Ernst 1953, Lewis and Lewis, 1955), and Smith-Huerta (1986) provided evidence from a small number of electrophoretically variable loci consistent with this interpretation. However, other possible progenitors were not examined, including *C. tembloriensis, C. exilis*, and *C. springvillensis*, which are very closely related to and almost morphologically identical to *C. unguiculata*. Our analysis supported *C. epilobioides* as one of the progenitors, but indicated that *C. exilis*, rather than *C. unguiculata*, was most likely the other progenitor.

*C. tenella* and *C. davyi* are morphologically similar tetraploids both with 17 chromosome pairs. Because of the ease of forming hybrids between the two species, as well as the high degree of chromosome pairing in those hybrids, it was speculated that they shared a common, unidentified tetraploid ancestor (Raven and Lewis 1959) derived from a species similar, but not identical, to one of the extant diploid species from Section *Godetia*. We infer that the most likely progenitors of both *C. davyi* and *C. tenella* were the diploids *C. speciosa* and *C. imbricata*, which are both extant members of Section *Godetia*. The relatively high PBH values for a third species, *C. williamsonii*, is likely due to the hybridization event occurring at RTU *≈* 0.3, which is soon after the split of this species and *C. imbricata* (RTU = 0.41). We also found that the estimated hybridization times were almost identical for the two tetraploid species, which is consistent with, but does not prove, the hypothesis that they share a tetraploid ancestor in common.

Based on morphological and cytological analyses (Ernst, 1953, Lewis and Lewis 1955), and a limited analysis of electrophoretic variants (Ford and Gottlieb 2003), it has been suggested that the diploid progenitors of *C. similis* were *C. modesta* and *C. epilobioides*. Unfortunately, *C. modesta* was one of the few diploid species we were unable to sample, so we could not directly evaluate this suggestion. However, our simulation results are consistent with this hypothesis, as one of the three best performing models simulates hybridization between *C. epilobiodes* and an unknown species that split from the progenitor of *C. biloba* and *C. lingulata* (i.e., split from branch 8→13 in Fig. 4A), which represents precisely what would be expected if the unknown species is *C. modesta*, as this species was found to be sister to *C. biloba* and *C. lingulata* in the phylogeny by Ford and Gottlieb (2003). Unfortunately, because we could not obtain specimens of *C. modesta* for our analyses, definitive confirmation needs further investigation.

Based on morphological and cytological evidence, Lewis and Lewis (1955) suggested that *C. rhomboidea* resulted from hybridization between *C. virgata* and *C. mildrediae*, and a limited analysis of electrophoretic variation is consistent with this suggestion (Smith-Huerta, 1986), though other possible progenitors were not examined. Our results do not support this suggestion, as simulations involving these two species produced a poor match to the observed data. The most likely progenitors are instead *C. stellata*, the sister species to *C. mildrediae* (Stanton et al., 2026), and one of two unknown species within Section *Myxocarpa*. Our simulations show that the most likely time of hybridization is between 0.01 and 0.2 RTU, which would make *C. rhomboidea* one of the most recently evolved *Clarkia* polyploids, and would also imply that the unknown species either is still extant, but undiscovered, or it went extinct fairly recently.

It has been previously recognized that *C. pulchella* must be an older allotetraploid (Lewis and Lewis 1955, Sytsma 1990), which is consistent with our results; no one single diploid received a majority of PBH from *C. pulchella* and all of the highest performing models involved hybridization between the ancestor of Section *Eucharidium* (branch R→26 in Fig. 6A) and another early lineage within *Clarkia*. Based on morphology, chromosome number, and phylogenetic analysis, it has been previously hypothesized that *C. pulchella* was likely formed by hybridization between a member of Section *Eucharidium* (likely *C. concinna* or its ancestor) and *C. virgata* or a closely related extinct species (Lewis and Lewis 1955; Sytsma et al. 1990), but our analyses seem to rule out these suggestions.

### Caveats and limitations of the method

One potential limitation of our approach is that it requires a strongly supported phylogenic tree that contains most, if not all, species that are potential diploid progenitors. In the case of *Clarkia* examined here, our phylogeny is highly supported and contains 28 of the 32 known diploid species. However, with larger genera, including all known species may be impractical. Nevertheless, it may instead be possible to use a subset of clades that contain the hypothesized progenitors to render the analysis more tractable.

In addition, it remains to be tested how well our method performs for progressively older allopolyploids, where biased fractionation between subgenomes and other widespread genomic alterations could progressively erode the homology between the progenitors and the polyploid (Cheng et al. 2018). The crown age of *Clarkia* has been roughly estimated in a larger analysis to be around 25 MYA (Freyman and Höhna 2019), so even the oldest allopolyploid tested in this analysis, *C. pulchella*, can be no older than this. Indeed, even in the case of *C. pulchella*, multiple models involving multiple potential ancestral and extinct progenitor species were equally as good fits for the data, and we were unable to conclusively determine the progenitor species for this polyploid, other than it likely did not involve any of the extant diploids in the genus.

Another possible limitation of our approach is that although we were able to assume a constant, identical substitution rate for all genes (molecular clock), it is well known that substitution rates can vary both over time and among genes (Bromham and Penny 2003, Lee et al. 2015). Evaluation of the extent to which relaxing our assumption would alter our conclusions is beyond the scope of this study. Nevertheless, our simulation model could be modified in a straightforward, though computationally more intensive, way to address this issue.

A final limitation of our approach is that it currently provides only limited statistical assessment of whether one pair of hypothesized progenitors provides a significantly better fit to the data than another pair. Our analysis accounts for variability in the Sum of Squares value due to running a finite number of simulations, and conservatively assumes that if the SS distributions for different models do not overlap, then the model with the lowest mean SS is the preferred model. However, this approach does not account for uncertainties in other variables such as estimated node ages in the phylogenetic tree or estimated substitution rates. Future improvements in our approach should be directed to addressing these limitations.

Despite these limitations, the methods presented here provide a quick and computationally light method for forming hypotheses about the progenitors for allopolyploid species, particularly in systems where no *a priori* evidence exists, that can then be refined for further, more extensive, testing and analysis.

## Acknowledgement

This work was supported by National Science Foundation (NSF) grant DEB 1542387 and by a sequencing voucher from Duke University Sequencing and Genomic Technologies. The authors would like to thank Dr. Norman Weeden at Montana State University, the California Botanical Garden, the University of California at Berkeley Botanical Garden, the US Department of Agriculture Seed Bank, and Dr. Kathleen Kay at University of California at Santa Cruz for providing the seeds that made this research possible.

## Author Contributions

K. Stanton: sample acquisition, species identification, sequencing, read mapping analysis, data interpretation, manuscript writing. M. Rausher: simulation analysis, funding, data interpretation, manuscript writing.

## Data Availability

RNA sequencing data is available in NCBI under BioProjects PRJNA1514240 and PRJNA1400954. Supplemental materials, such as newick format tree file, photographs of all polyploid plant lines, transcriptome assemblies, and the proportion best hit data, are all available on Dryad (doi.org/10.5061/dryad.m0cfxppkp). Code used for the simulations will be uploaded on GitHub.

## References

Bird, K.A., VanBuren, R., Puzey, J.R. and P. Edger. 2018. The causes and consequences of subgenome dominance in hybrids and recent polyploids. New Phytologist 220: 87–93. 10.1111/nph.15256

Blanc, G., and K.H. Wolfe. 2004. Widespread paleopolyploidy in model plant species inferred from age distributions of duplicate genes. The Plant Cell 16: 1667–1678.

Blischak, P.D., C.E. Thompson, E.M. Waight, L.S. Kubatko, and A.D. Wolfe. 2020. Inferring Patterns of Hybridization and Polyploidy in the Plant Genus Penstemon (Plantaginaceae). Biorxiv doi: 10.1101/2020.09.04.283093.

Bromham, L., and D. Penny. 2003. The modern molecular clock. Nature Reviews Genetics 4: 216–224.

Cheng, F., Wu, J., Cai, X., Liang, J., Freeling, M., and X. Wang. 2018. Gene retention, fractionation and subgenome differences in polyploid plants. Nature plants 4:258–268.

Cui, L., P.K. Wall, J.H. Leebens-Mack, B.G. Lindsay, D.E. Soltis, J.J. Doyle,… and C.W. dePamphlis. 2006. Widespread genome duplications throughout the history of flowering plants. Genome Research. 16:738–749.

Doyle, J.J., and A.N. Egan. 2010. Dating the origins of polyploidy events. New Phytologist. 186: 73–85.

Ernst, W. 1953. The derivation of Clarkia delicata and Clarkia similis, two allotetraploid species. M. A. Thesis, University of California, Los Angeles.

Freyman, W.A., and S. Höhna. 2019. Stochastic character mapping of state-dependent diversification reveals the tempo of evolutionary decline in self-compatible Onagraceae lineages. Systematic Biology 68:505–519.

Ford, V.S., and L.D. Gottlieb. 2003. Reassessment of the phylogenetic relationships in Clarkia sect. Sympherica. American Journal of Botany 90: 284–292.

Grabherr, M.G., Haas, B.J., Yassour, M., Levin, J.Z., Thompson, D.A., Amit, I., … and A. Regev. 2011. Trinity: reconstructing a full-length transcriptome without a genome from RNA-Seq data. Nature biotechnology 29:644.

Hurka, H., N. Friesen, D.A. German, A. Franzke, and B. Neuffer. 2012. Missing link’ species Capsella orientalis and Capsella thracica evolution of model plant genus Capsella (Brassicaceae). Molecular Ecology 21: 1223–1238.

Kaderei, J.W., S. Uribe-Convers 1, E. Westberg, and H.P. Comes. 2006. Reciprocal hybridization at different times between Senecio flavus and Senecio glaucus gave rise to two polyploid species in north Africa and south-west Asia. New Phytologist 169: 431–441.

Kagale, S., S.J. Robinson, J. Nixon, R. Xiao, T. Huebert, J. Condie,… and A.P. Parkin. 2014. Polyploid Evolution of the Brassicaceae during the Cenozoic Era. The Plant Cell. 26: 2777–2791.

Kozlov, A.M., D. Darriba, T. Flouri, B. Morel, and A. Stamatakis. 2019. RAxML-NG: a fast, scalable and user-friendly tool for maximum likelihood phylogenetic inference. Bioinformatics 35: 4453–4455.

Lee H.J., N. Rodrigue, and J.L. Thorne. 2015. Relaxing the molecular clock to different Degrees for different Substitution types. Mol Biol Evol. 32:1948–61.

Lewis, H. and M.E. Lewis. 1955. The genus Clarkia. University of California Publications in Botany 20: 241–392.

Paradis, E., Claude, J., and K. Strimmer, 2004. APE: Analyses of Phylogenetics and Evolution in R language. Bioinformatics 20:289–290. 10.1093/bioinformatics/btg412

Patro, R., Duggal, G., Love, M.I., Irizarry, R.A., and C. Kingsford. 2017. Salmon provides fast and bias-aware quantification of transcript expression. Nature methods 14: 417–419.

Rana, D., T. van den Boogaart, C.M. O’Neill, L. Hynes, E. Bent, L. Macpherson, J.Y Park, Y.P. Lim, and I. Bancroft. 2004. Conservation of the microstructure of genome segments in Brassica napus and its diploid relatives. The Plant Journal 40: 725–733.

Raven, P.H., and H. Lewis. 1959. The relationship of clarkias from two continents. Brittonia 11:193–205.

Sianta, S.A., and K.M. Kay. 2022. Phylogenomic analysis does not support a classic but controversial hypothesis of progenitor-derivative origins for the serpentine endemic Clarkia franciscana. Evolution 76:1246–1259.

Smith-Huerta, N.L. 1986. Isozymic diversity in three allotetraploid Clarkia species and their putative diploid progenitors. Journal of Heredity 77:349–354.

Solis-Lemus, C., and C. Ane. 2016. Inferring phylogenetic networks with maximum pseudolikelihood under incomplete lineage sorting. PLoS Genetics: 12: 3005896.

Solís-Lemus, C., P. Bastide, and C. Ané. 2017 PhyloNetworks: A Package for Phylogenetic Networks. Mol Biol Evol. 34:3292–3298.

Stanton, K., Zarei, M., and M. Rausher. 2026 Phylotranscriptomics resolves species-level phylogeny for the genus Clarkia (Onagraceae). Preprint: 10.2139/ssrn.5954958

Sytsma, K., J.F. Smith, and L.D. Gottlieb. 1990. Phylogenetics in Clarkia (Onagraceae): restriction site mapping of chloroplast DNA. Systematic Botany 15: 280–295.

Wagner, W.L., Hoch, P.C., and P.H. Raven. 2007. Systematic botany monographs: revised classification of the Onagraceae. Systematic Botany Monographs, 83:1–243.

Wang, N., L.J. Kelly, H.A. McAllister, J. Zohren, and R.J.A. Buggs. 2021. Resolving phylogeny and polyploid parentage using genus-wide genome-wide sequence data from birch trees. Molecular Phylogenetics and Evolution 160: 107126.

Wood, T.E., N. Takebayashi, M.S. Barker, I. Mayrose, P.B. Greenspoon and L.H. Rieseberg. 2009. The frequency of polyploid speciation in vascular plants. Proc. Nat. Acad. Sci. 106: 13875–13879.

Yan, Z., Cao, Z., Liu, Y., Ogilvie, H. A., and L. Nakhleh. 2022. Maximum parsimony inference of phylogenetic networks in the presence of polyploid complexes. Systematic Biology 71:706–720.

Yan, Z., Cao, Z., and L. Nakhleh. 2024. Polyphest: fast polyploid phylogeny estimation. Bioinformatics 40:ii20–ii28.

Zhang, C., M. Rabiee, E. Sayyari, and S. Mirarab. 2018. ASTRAL-III: Polynomial time species tree reconstruction from partially resolved gene trees. BMC Bioinformatics 19: 15–30.

Zohary, C., J.R. Harlan, and A. Vardi. 1969. The wild diploid progenitors of wheat and their breeding value. Euphytica 18: 58–65.

Zou, X-H., Y-S Du, L. Tang, X-W Xu, J.J. Doyle, T. Sang, and S. Ge. 2015. Multiple origins of BBCC allopolyploid species in the rice genus (Oryza). Scientific Reports 5:14876.

