## Supplemental Materials for "Resolving Allopolyploid Origins Within the Genus *Clarkia* Using a Novel Read-Mapping and Modeling Approach"

### Supplementary Material

#### I. Phylogenetic Trees

##### *A. Comparison of ultrametric trees*

We constructed ultrametric trees based on our RAxML tree using chronos from the R *ape* package (Paradis et al 2004) under various assumptions about a molecular clock. These trees were then to identify the "best" tree by comparing both likelihood scores and the Penalized Hierarchical Integrated Information Criterion (PHIIC). The latter is similar to the Akaike Information Criterion (AIC), in that the tree with the lowest score is considered the best tree among the alternatives.

Among criterion tested, all trees yielded very similar likelihoods. However, the PHIIC was substantially lower for the strict-clock tree (see table below), indicating there is no reason to reject the assumption of a strict clock. Under the strict clock, the substitution rate is estimated to be 0.1913893 substitutions per site per 1 relative time unit (RTU), where 1 RTU is the time from the root to the tip of the tree.

| Clock Model | λ parameter | log-likelihood | PHIIC |
|---|---|---|---|
| Correlated | 1 | -9.3329 | 178.67 |
| Correlated | 100 | -9.3417 | 178.68 |
| Correlated | 1000 | -9.3440 | 178.69 |
| Relaxed | 1 | -9.3228 | 178.65 |
| Relaxed | 10 | -9.3797 | 178.77 |
| Relaxed | 100 | -9.3844 | 178.81 |
| Discrete* | NA | -9.3443 | 108.69 |
| Strict | NA | -9.3443 | 72.69 |

* 10 rate bins

#### *B. Ultrametric tree used in simulations*

The ultrametric tree, branch lengths, and expected number of substitutions per site per branch are shown in Fig. SI.1 and SI.2. Expected substitutions per site per branch are calculated as branch length (in RTU) x 0.1913893.

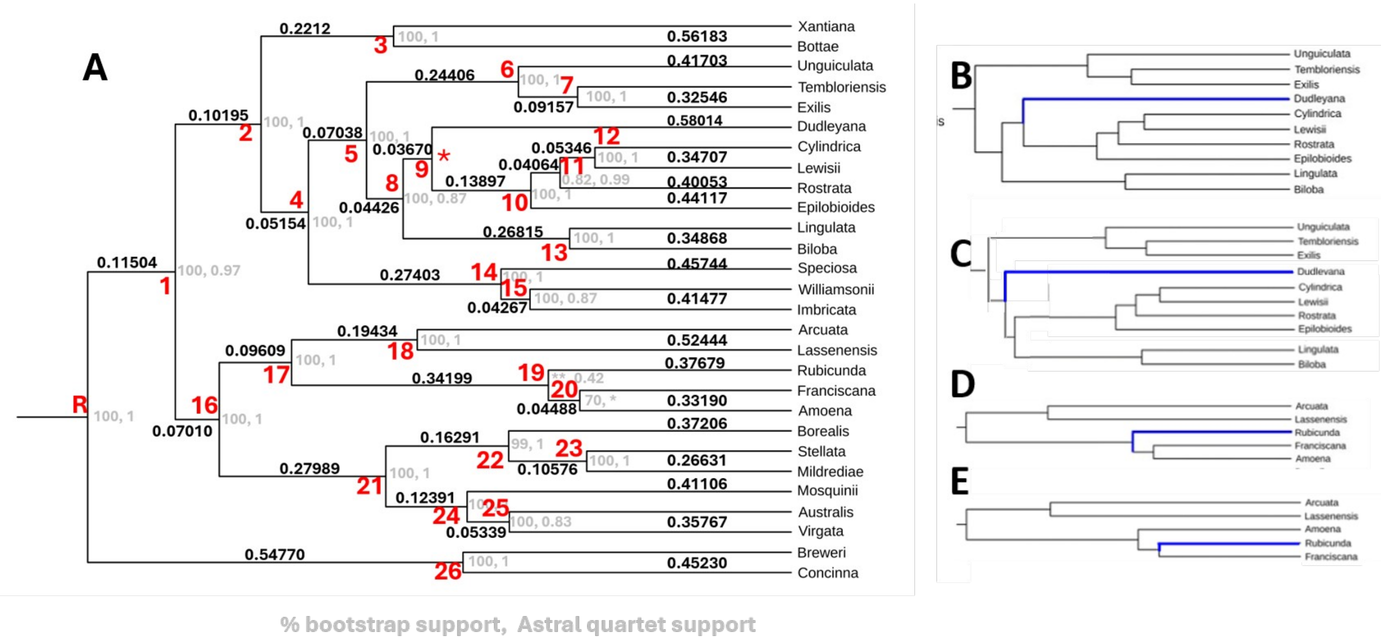


**Figure SI.1**. *Clarkia* phylogeny adapted from Stanton et al. 2026. **A.** Ultrametric tree corresponding to RAxML tree of Stanton et al. (2026). Black numbers are relative branch lengths. Grey numbers indicate node relative support. First number is % of Bootstrap runs that support node. Second number is Astral quartet support. Grey asterisks indicate nodes where there is disagreement between the RAxML and Astral reconstructions. Red asterisk indicates node corresponding to disagreement between the two reconstructions over placement of *C. dudleyana.* **B. and C.** Alternative placements of *C. dudleyana* (blue branch) for RAxML tree (B.) and Astral tree (C.). **D. and E.** Alternative relationships among *C. rubicunda* (blue branch), *C. franciscana*, and *C. amoena* for RAxML tree (D.) and Astral tree (E.).

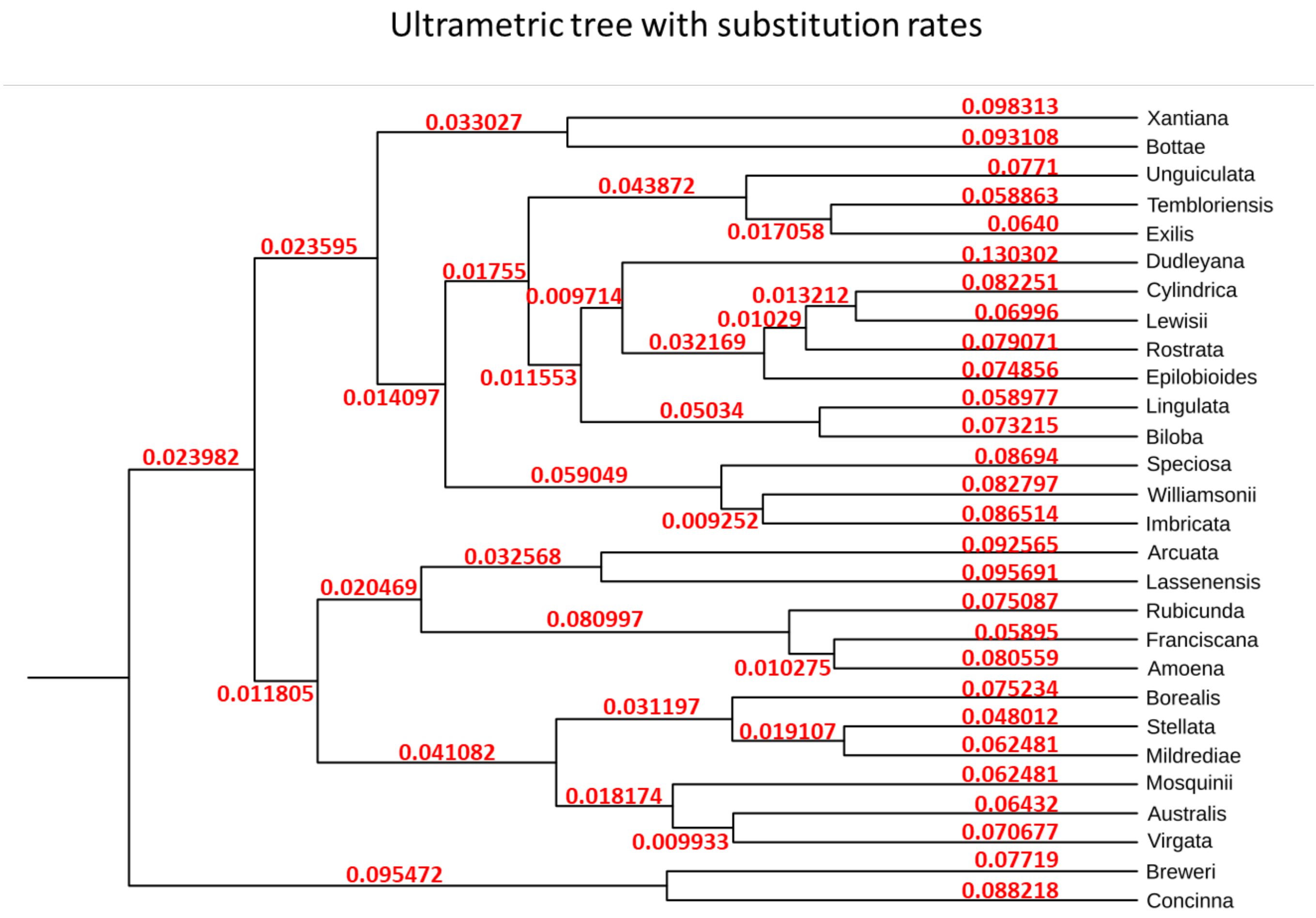


**Figure SI.2.** Ultrametric tree corresponding to RAxML tree of Stanton et al. (2026). Numbers on branches are estimated substitution rates (number of substitutions per nucleotide site along branch).

#### II. Simulation Methods

The purpose of the simulations is to predict the similarity between diploid species genomes and the genomes of two hypothesized tetraploid progenitor species. Similarity is measured as proportion of "best hits" of simulated RNA reads (see below). Each simulation is based on an assumed phylogeny (ultrametric tree, Fig SI.2), an assumed substitution rate λ (see Supplementary Material SI), an assumed pair of diploid progenitors, an assumed time of hybridization, and an assumed sub-genome dominance factor. For each pair of assumed diploid progenitors, multiple possible hybridization times were examined in different simulations. For a detailed explanation of the simulation protocol, we refer to the hypothetical phylogeny depicted in Fig. SII.1.

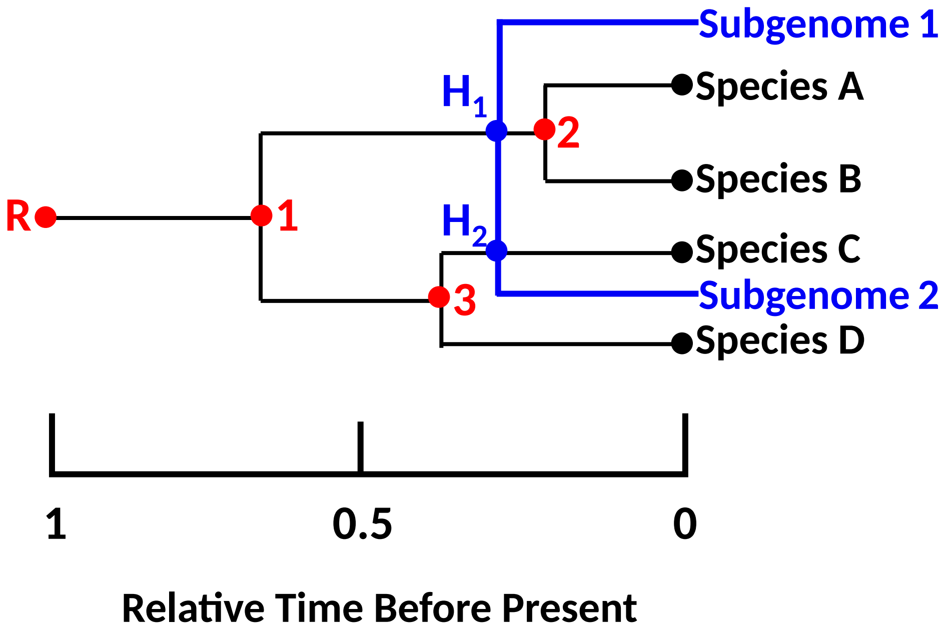


**Figure SII.1.** Hypothetical phylogenetic tree used to illustrate method of simulating evolution. Red circles: internal nodes. Black circles: terminal nodes (diploid species). Blue circles: Hybridization nodes. Blue lines: lineages of hybridizing subgenomes of tetraploid. Times of nodes are represented in relative time units ("RTUs"), where 1 RTU is the time from the root to the tip of the tree.

##### Simulation Algorithm

Each simulation consists of a series of steps. We illustrate these steps for the hypothetical phylogeny depicted in Fig. SII.1. For each assumed hybridization time we ran 1,000 replicate simulations. The steps for each replicate are as follows:

**1. Establish the Root genome.**

Genomes are represented by a vector of length 30,000 bp, $\boldsymbol{G_{rk}}$. The subscript $r$ refers to the replicate number and the subscript $k$ indicates the genome at node $k$ of the phylogeny. Each element $i$ of the vector, $\boldsymbol{g_{rk}(i)}$, corresponds to a nucleotide, and each consecutive 150 bp segment represents a "gene". Each element $\boldsymbol{g_{rk}(i)}$can take on any of 4 values (0, 1, 2, or 3) representing the four possible nucleotides. The genome of the tree root (node R), $\boldsymbol{Gr_R}$, consists of all 0's, i.e. $\boldsymbol{g_{rR}(i)} = 0$ for all $i \in [1, \ldots, 30{,}000]$.

Genomes are divided into 40 contiguous "genes", each gene consisting of 750 nucleotides.

**2. Simulate evolution along branch node R→ node 1 (abbreviated as $\boldsymbol{B_{R\to 1}}$).**

The length of this branch is $\boldsymbol{\tau_{R\to 1}} = \boldsymbol{T_R - T_1}$, where $\boldsymbol{T_i}$ is the time in RTU's (see Fig. SII.1) of node $i$. The expected number of substitutions at each site along this branch is, $\boldsymbol{S_{R\to 1}}$, is

$$\boldsymbol{S_{R\to 1} = \tau_{R\to 1}\ \lambda}$$

For each site, a random number is drawn from a Poisson distribution with mean $\boldsymbol{S_{R\to 1}}$ to determine that number of substitutions at that site along the branch.. For most sites, this number is either 0 or 1, with a small proportion of sites having 2 changes. Of those having 2 changes, 2/3 will have a nucleotide different from the original nucleotide. The total number of sites changing along the branch is thus taken to be $N_{sub}$ = (number of sites with 1 change) + 2/2 (number of sites with 2 changes). At each of these sites, the original nucleotide in $\boldsymbol{G_{rR}}$ is changed randomly to one of the three other nucleotides, producing a new genome vector, $\boldsymbol{G_{r1}}$.

**3. Simulate evolution along branches $\boldsymbol{B_{1\to 3}}$ and $\boldsymbol{B_{3\to Species\ D}}$.**

Changes along these branches are simulated in similar fashion, with the initial genomes corresponding to $\boldsymbol{G_{r1}}$ and $\boldsymbol{G_{r3}}$ and branch lengths $\square_{1\to 3}=T_1-T_3$ and $\square_{3\to Species\,D}=T_3$, respectively, to yield genomes $\boldsymbol{G_{r3}}$ and $\boldsymbol{G_{r\ Sprecies\ D}}$.

**4. Simulate evolution along branch $B_{1\to H_1\square}$.**

Changes along this branch are simulated in similar fashion, with the initial genome $\boldsymbol{G_{r1}}$ and $\square_{1\to H_1}=T_1-T_{H_1}$, to yield the genome at node $\mathbf{H_1}$, $G_{r\,H_1}$

**5. Simulate evolution of tetraploid Subgenome 1 along branch $B_{H_1\to Subgenome\,1\square}$.**

Changes along this branch are simulated in similar fashion, with the initial genome $G_{r\,H_1}$ and $\square_{H_1\to Subgenome\,1}=T_{H_1}$, to yield the current Subgenome 1 of the tetraploid, $G_{r\,Subgenome\,1}$.

6. **Simulate evolution of tetraploid Subgenome 2 along branch** $B_{H_2 \rightarrow Subgenome 2}$❑.

Changes along this branch are simulated in similar fashion, with the initial genome $\boldsymbol{G_{r2}}$, and ❑$_{H_2 \rightarrow Subgenome 2} = T_{H_2} = T_{H_1}$, to yield the current Subgenome 1 of the tetraploid, $G_{r\,Subgenome 2}$.

7. **Simulate evolution along remaining branches.**

Changes along the remaining branches ($B_{H_1 \rightarrow 2}, B_{2 \rightarrow Species A}, B_{2 \rightarrow Species B}, B_{H_2 \rightarrow Species C}$),were simulated in similar manner to produce the genomes of the extant diploid species ($\boldsymbol{G_{r\,Species\,A}}$, $\boldsymbol{G_{r\,Species\,B}}$, and $\boldsymbol{G_{r\,Species\,C}}$)

8. **Calculate simulated Proportion Hits to each diploid species**

Proportion hits of each subgenome were determined by assessing the similarity between each subgenome and each diploid species genome.

Let $\boldsymbol{R_{r1ij}}$ be "read" *j* of from "gene" *i* of the tetraploid sub-genome $\boldsymbol{G_{r1}}$, where $j \in \{1,2,3,4,5\}$. We define a "read" as one of 5 contiguous 150 bp segments of a particular gene in the subgenome. In these simulations, a "read" corresponds to an RNA read. In particular

$R_{r1ij} = G_{r1}[\phi_{ij}+1, \phi_{ij}+2, \ldots, \phi_{ij}+150]$,

where $\phi_{ij} = (i-1)750 + (j-1)150$.

For example, for read 1 from gene 1, $\phi_{11} = (1-1)750 + (1-1)150 = 0$ and $R_{r111} = G_{r1}[1,2,\ldots,150]$. Similarly, for read 2 from gene 5, $\phi_{11} = (5-1)750 + (2-1)150 = 3150$ and $R_{r152} = G_{r1}[3151, 3152, \ldots, 3300]$.

Similarly, define "target" $\boldsymbol{T_{rkij}}$ to be the 150 bp segment of diploid genome *k* corresponding to the position of read $\boldsymbol{R_{r1ij}}$ in subgnome 1.

Next, define the divergence between sequence of read *j* of gene *i* from subgenome 1 and the sequence of the target of read *j* of gene *i* from diploid genome *k*, $\boldsymbol{D_{r1kij}}$ , to be the number of nucleotides of that are different between the two sequences, i.e. the Hamming Distance (XXX) between the reads:

$$\boldsymbol{D_{r1kij}} = Ꞣ(\boldsymbol{R_{r1ij}}, \boldsymbol{T_{rkij}})$$

where Ꞣ$(x,y)$ is the Hamming Distance between vectors *x* and *y*. Define the vector $\boldsymbol{D_{r1\bullet j}}$ as the vector of Hamming Distances for Subgenome 1:

$\boldsymbol{D}_{r1\bullet j} = \{ \boldsymbol{D}_{r11ij}, \boldsymbol{D}_{r12ij}, \ldots, \boldsymbol{D}_{r1kij}, \ldots \}$

Finally, let $\boldsymbol{h}_{r1\bullet ij}$ be a vector of "best matches" or "hits" for "read" *j* of gene *i* between Subgenome 1 and the genomes of each of the *n* diploid species, i.e.

$$\boldsymbol{h}_{r1\bullet ij} = \{\eta_1, \eta_2, \ldots, \eta_n\}$$

$$\text{where } \square_k = \begin{cases} 0 \; if \; D_{r1kij} > \min_k D_{r1kij} \\ \frac{1}{m} \; if \; D_{r1kij} = \min_k D_{r1kij} \end{cases}$$

Here, *m* is the number of species for which the distance equals the minimum distance. Because in real genomes, it is very unlikely that two or more diploids will be best matches, if m > 1, we randomly assign one such diploid to have $\square_k = 1$and the other(s) to have $\square_k = 0$. This procedure produces a vector $\boldsymbol{h}_{r1\bullet ij}$ with all zeros except a 1 in the position corresponding to the diploid species with the best hit. This procedure is then repeated with the remaining reads of gene *i*.

Next, we calculate, for each gene, whether gene *i* in diploid species *k* has has a "best hit" from any read from gene *i* in subgenome 1. This produces a vector $\boldsymbol{H}_{r1\bullet i\bullet}$ which has *n* elements, each of which is either 0 if there are no best hits for a given diploid species, or a 1 if there are best hits for that species. This step corresponds to counting best hits for a gene in a diploid species only as present or absent when calculating the observed number of best hits from transcriptome reads. In similar fashion, a vector of best hits from subgenome 2 for gene *i*, $\boldsymbol{H}_{r2\bullet i\bullet}$, is calculated.

If there are no dominance effects, then for gene *i* a combination of $\boldsymbol{H}_{r1\bullet i\bullet}$ and $\boldsymbol{H}_{r2\bullet i\bullet}$ would represent all the best hits from both subgenomes. This vector, $\boldsymbol{H}_{r\bullet\bullet i\bullet}$ would have elements

$$H_{rki} = \begin{cases} 1 \; if \; H_{r1i}[k] = 1 \lor H_{r2i}[k] = 1 \\ 0 \; othewise \end{cases} \qquad \text{Eq. SII.1}$$

Finally, a vector of proportion of best hits from both subgenomes is calculated as

$$\boldsymbol{p}_r = \frac{\sum_i H_{ri}}{(\sum_i H_{ri}) \circ \vec{1}} \qquad \text{Eq. SII.2}$$

where $\circ$ indicates an inner (dot) product, and $\vec{1}$ is a vector of 1's of length *n*, the number of diploid species.

Subgenome dominance, in which genes from one subgenome are more highly expressed or are less likely to be inactivated or deleted, is common in plants (XXX). To allow for this, we introduced an additional factor in our simulations. We let the parameter ***D***, which we allow to take on values -0.3, -0.2, -0.1, 0, 0.1, 0.2, or 0.3, account for such dominance. When ***D*** = 0, the procedure above, with best hits from the two genome combined using Eq. SII.1, is appropriate. However, for other values of ***D***, we adopt a different approach. Specifically, we let $\Delta=|D|$ be the probability that best hits for a particular gene from a particular subgenome is not included in $\boldsymbol{H}_{r\bullet\bullet i\bullet}$. Specifically, if ***D*** < 0, Eq. SII.1 is modified to

$$H_{rki}=\begin{cases} 1\, if\ rand>\Delta \wedge (H_{r1i}[k]=1 \vee H_{r2i}[k]=1 \\ 1\, if\ rand\leq\Delta \wedge H_{r2i}[k]=1 \\ 0\, otherwise \end{cases} \quad \text{Eq. SII.3}$$

where *rand* is a random number between 0 and 1. This essentially treats a fraction **Δ** of genes as absent or not expressed in subgenome 1.

By contrast, if ***D*** > 0, Eq. SII.1 is modified to

$$H_{rki}=\begin{cases} 1\, if\ rand>\Delta \wedge (H_{r1i}[k]=1 \vee H_{r2i}[k]=1 \\ 1\, if\ rand\leq\Delta \wedge H_{r1i}[k]=1 \\ 0\, otherwise \end{cases} \quad \text{Eq. SII.4}$$

which treats a fraction Δ of genes as absent or not expressed in subgenome 2.

In our simulations we used different values of ***D*** ranging from −0.3 to 0.3. We base this range on values of ***D*** taken or estimated from the literature (Table SII.1), all of which have an absolute value of 0.3 or less. For most of these estimates, the dates of hybridization are in the tens of millions of years. Because most species of *Clarkia* are believed to have originated <10 MYA (Freyman et al., 2019), we believe that this value is a reasonable upper bound to use in our simulations.

**Table SII.1.** Estimates of sub-genome dominance in tetraploids taken or calculated from the literature. Dominance is 1 – the relative retention of genes in the two subgenomes of a tetraploid.

| **Species** | **Age (my)** | **Dominance (*D*)** | **Data Source** | **Calculation** | **Reference** |
|---|---|---|---|---|---|
| *Heritiera littoralis* | 60 | 0.08974 | text P. 6 | 1 | Feng et al. in press |
| *Zea mays* | 5 −12 | 0.31576 | Fig. 2 | 2 | Schnable et al. 2011 |
| *Biscutella laevigata* | 11 | 0.29412 | Fig. 2a | 3 | Beringer et al. 2024 |
| *Arabidopsis* | 25-40 | 0.214 | Table 1 | 4 | Emery et al. 2018 |

*thaliana*

**Calculations**

subgenome 1: 5121 single copy genes; subgenome 2: 4156 single copy genes; 5627 two-copy
1 genes
Dom = 1-(4156+5627)/(5121+5627) =
0.089784

2 1-(avg. retention on subgenome 2)/(avg. retention on subgenome 1)

subgenome LF: 5000 single copy genes; subgenome MF: 3500 single copy genes; 3500 two-copy
3 genes
Dom = 1-(2500+3500)/(5000+3500) = 0.29412

4 1 - bias strength for At-α, High-synteny: optimized order

Example of calculations for Step 8

As an illustration of these calculations, consider an analysis of a genome consisting of 3 genes (Fig. SII.2). Read 1 from gene 1 of subgenome 1 ($\boldsymbol{R}_{r111}$) is a best match for the corresponding target sequence in diploid species 1 ($\boldsymbol{T}_{r111}$), as indicated by the left-most red dots and bar in Fig. SII.2. This means that $\boldsymbol{h}_{r1\bullet 11} = \{1, 0, 0, 0, 0\}$, i.e. the first element of the vector is 1, indicating the best hit is to species 1. By contrast, reads 2-5 from gene 1 from subgenome 1 are best matches for the corresponding target sequences in diploid species 2, as indicated by the 2nd – 5th red dots and bar. This means that $\boldsymbol{h}_{r1\bullet 1j} = \{0, 1, 0, 0, 0\}$ for $j \in \{2,3,4,5\}$, indicating the best matches of these reads are to diploid species 2 because the "1" is the second element of these vectors. Consequently, for gene 1, there are only two diploid species that have best hits from subgenome 1 reads: species 1 and 2. In turn, this means that the vector of best hits for gene 1 is $\boldsymbol{H}_{r1\bullet 1\bullet} = \{1, 1, 0, 0, 0\}$.

In similar fashion, read 2 from gene 1 of subgenome 2 (($\boldsymbol{R}_{r212}$), is a best match for the corresponding target sequence in diploid species 5 ($\boldsymbol{T}_{r512}$), as indicated by the blue dots and bar second from the left. This means that $\boldsymbol{h}_{r2\bullet 12} = \{0, 0, 0, 0, 1\}$, i.e. the 5th element of the vector is 1, indicating the best hit is to diploid species 5. By contrast, reads 1 and 3-5 from gene 1 from subgenome 2 are best matches for the corresponding target sequences in diploid species 4, as indicated by the blue dots and bar that are first and 3rd – 5th from the left. This means that $\boldsymbol{h}_{r2\bullet 1j} = \{0, 0, 0, 1, 0\}$ for $j \in \{1,3,4,5\}$, indicating the best matches of these reads are to diploid species 4 because the "1" is the 4th element of these vectors. Consequently, for gene 1, there are only two diploid species that have best hits from subgenome 2 reads: species 4 and 5. In turn, this means that the vector of best hits for gene 1 is $\boldsymbol{H}_{r2\bullet 1\bullet} = \{0, 0, 0, 1, 1\}$.

Assuming no subgenome dominance, there are best hits for gene 1 from subgenomes 1 and 2 together to four diploid species: 1, 2, 4, and 5. Therefore, the best hit vector for gene 1 is $\boldsymbol{H}_{r\bullet\bullet 1\bullet} = \{1, 1, 0, 1, 1\}$. Another way of interpreting this is that $\boldsymbol{H}_{r\bullet\bullet 1\bullet}$ is a vector in which each element $k$ is 1 if there is one or more red or blue dots for Gene 1 in diploid species $k$ in Fig. SII.2 and 0 otherwise.

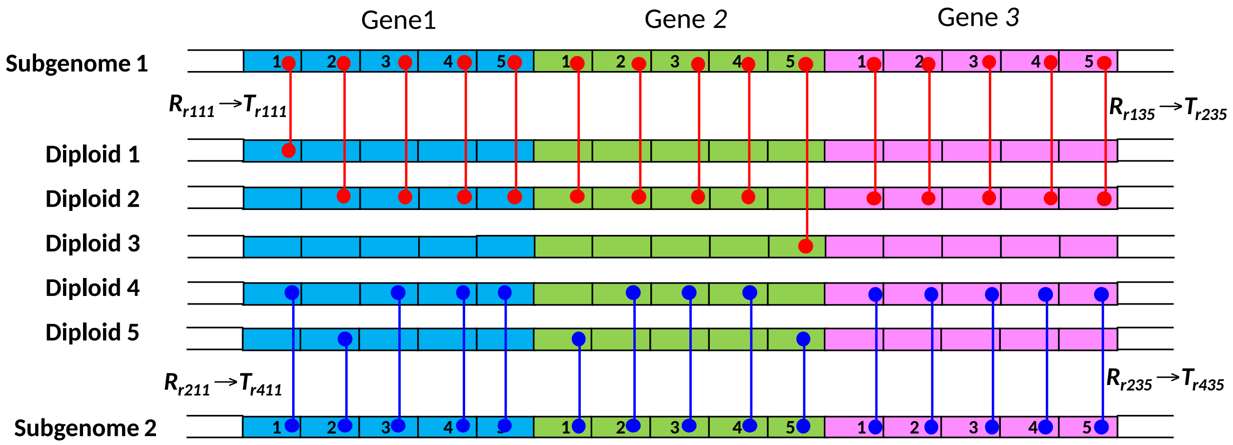


$h_{r1\bullet 11} = \{1,0,0,0,0\}$ $h_{r1\bullet 21} = \{0,1,0,0,0\}$ $h_{r1\bullet 31} = \{0,1,0,0,0\}$

$h_{r1\bullet 12} = \{0,1,0,0,0\}$ $h_{r1\bullet 22} = \{0,1,0,0,0\}$ $h_{r1\bullet 32} = \{0,1,0,0,0\}$

$h_{r1\bullet 13} = \{0,1,0,0,0\}$ $h_{r1\bullet 23} = \{0,1,0,0,0\}$ $h_{r1\bullet 33} = \{0,1,0,0,0\}$

$h_{r1\bullet 14} = \{0,1,0,0,0\}$ $h_{r1\bullet 24} = \{0,1,0,0,0\}$ $h_{r1\bullet 34} = \{0,1,0,0,0\}$

$h_{r1\bullet 15} = \{0,1,0,0,0\}$ $h_{r1\bullet 25} = \{0,0,1,0,0\}$ $h_{r1\bullet 35} = \{0,1,0,0,0\}$

$H_{r1\bullet 1\bullet} = \{1,1,0,0,0\}$ $H_{r1\bullet 2\bullet} = \{0,1,1,0,0\}$ $H_{r1\bullet 3\bullet} = \{0,1,0,0,0\}$

$H_{r2\bullet 1\bullet} = \{0,0,0,1,1\}$ $H_{r2\bullet 2\bullet} = \{0,0,0,1,1\}$ $H_{r2\bullet 3\bullet} = \{0,0,0,1,0\}$

$P_r = \{0.1, 0.3, 0.1, 0.3, 0.2\}$ $H_{r\bullet\bullet 1\bullet} = \{1,1,0,1,1\}$ $H_{r\bullet\bullet 2\bullet} = \{0,1,1,1,1\}$ $H_{r\bullet\bullet 3\bullet} = \{0,1,0,1,0\}$

**Figure SII.2. Hypothetical example of calculations for a genome with 3 genes, 5 "reads" per gene, and 5 diploid species. Genes are represented by different colors. "Reads" within genes are represented by the numbers 1, 2 . . . 5. Red dots connected by red line indicate the best match of a subgenome 1 read $R_{r1ij}$ to a corresponding diploid genome target $T_{rkij.}$. Blue dots connected by blue line indicate the best match of a subgenome 2 read $R_{r2ij}$ to a corresponding diploid genome target $T_{rkij..}$ Labels adjacent to red and blue lines indicate examples of read and corresponding target labels. Symbols for vectors are defined in text. Subscript "*r*" in all vectors indicates simulation replicate *r*.**

As illustrated in Fig. SII.2, the best hit vectors for genes 2 and 3 are calculated in similar fashion. For gene 2, $\boldsymbol{H}_{r\bullet\bullet 2\bullet} = \{0, 1, 1, 1, 1\}$ because subgenome 1 has best hits to species 2 and 3 and subgenome 2 has best hits to species 4 and 5; and for gene 3, $\boldsymbol{H}_{r \bullet \bullet 3 \bullet}$ = {0, 1, 0, 1, 0} because subgenome 1 has best hits to only species 2 and subgenome 2 has best hits to only species 4.

Finally, the proportion of best hits to each diploid species is, from Eq. SII.2,

$$P_r = \frac{\{1,1,0,1,1\}+\{0,1,1,1,1\}+\{0,1,0,1,0\}}{(\{1,1,0,1,1\}+\{0,1,1,1,1\}+\{0,1,0,1,0\}) \circ \{1,1,1,1\}}$$

$$\frac{\{1,3,1,3,2\}}{10}$$

$$\{0.1, 0.3, 0.1, 0.3, 0.2\}$$

**9. Repeat steps 1. – 8. for each replicate**

**10. Calculate average proportions of best hits, $\overline{P_r}$ across replicates and standard error.**

**11. Calculate Sum of Squares**

We use the sum of the squared deviation between $\overline{P_r}$ and the observed proportion best hits, $\boldsymbol{P}_{obs}$, as a measure of the lack of fit of the simulation to the observed data.

The sum of squares is calculated as

$$SS = \sum_{k=1}^{n} \left( \overline{P_r}[k] - P_{obs}[k] \right)^2$$

where $n$ is the number of diploid species and the $k$ in brackets designates the $k^{th}$ element corresponding to diploid species $k$.

#### III. Supplementary Figures

### *C. delicata*

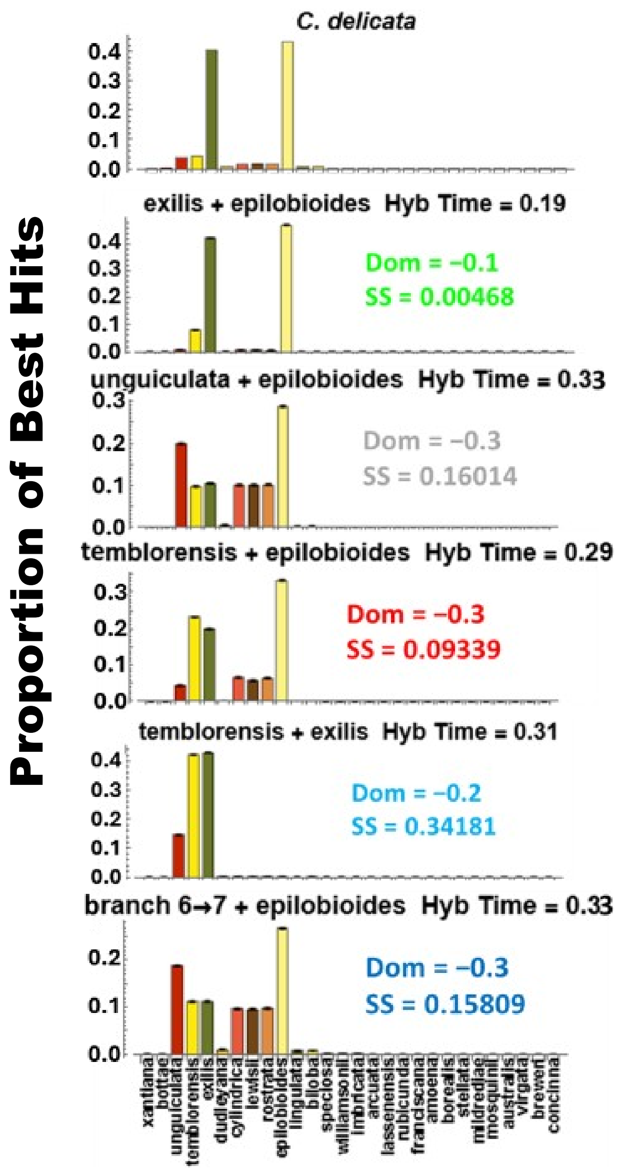


**Figure SIII.1.** Observed PBH spectrum for *C. delicata* (top), compared to simulated PBH spectra for each pair of possible progenitors examined. “Dom” is the value of subgenome dominance for the model. SS is Sum of Squared deviations for the model.

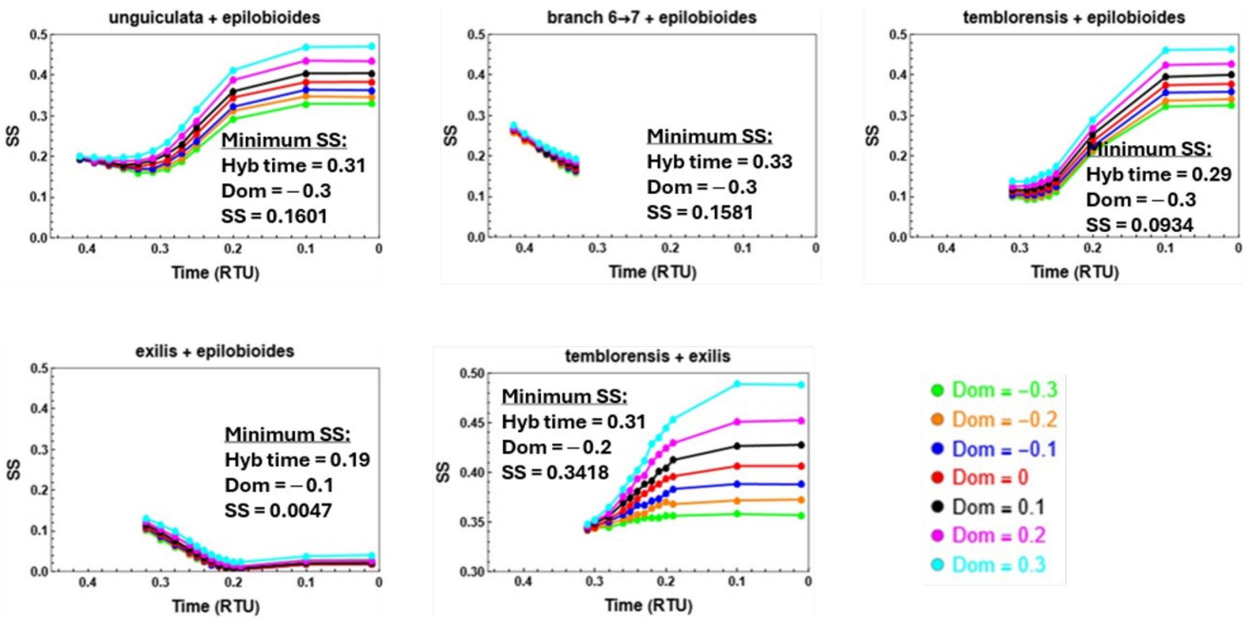


**Figure SIII.2.** Sums of Squared deviations (SS) between observed PBH spectra and predicted PBH for simulations of hybridizations that produced the tetraploid *C. delicata.* Each panel represents a different pair of hybridizing lineages (species) listed above each graph. Within each panel, each curve corresponds to different assumed sub-genome dominance values ("Dom")—see key at bottom right. X-axis in each part corresponds to different possible hybridization times before present, expressed as RTU.

### *C. davyi*

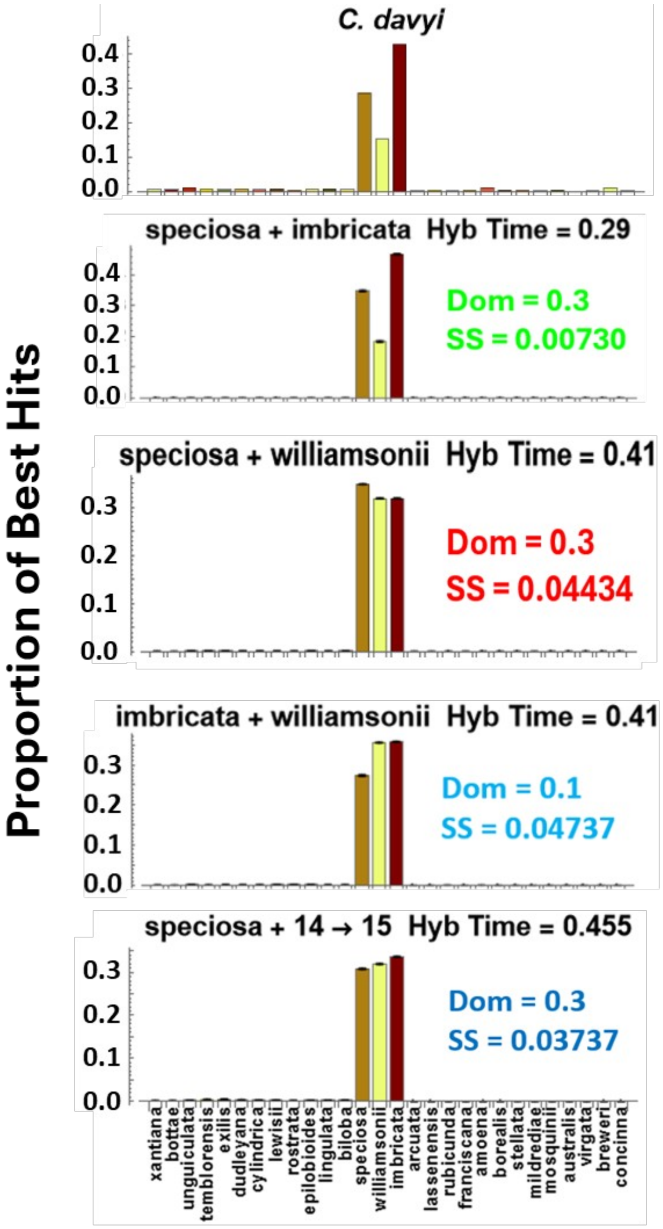


**Figure SIII.3**. Observed PBH spectrum for *C. davyi,* compared to simulated PBH spectra for each pair of possible progenitors examined. “Dom” is the value of subgenome dominance for the model. SS is Sum of Squared deviations for the model.

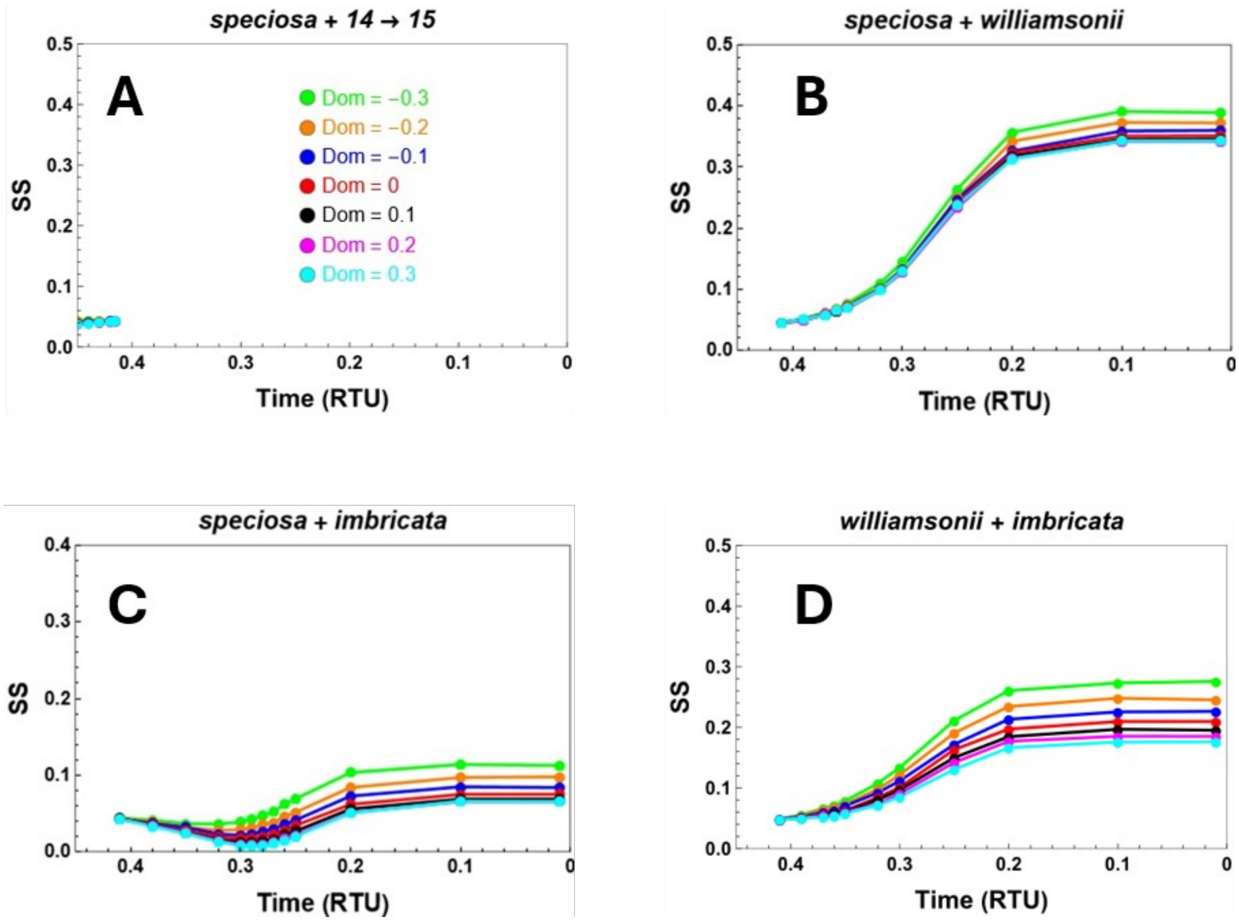


**Figure SIII.4**. Sums of Squared deviations (SS) between observed PBH spectra and predicted PBH for simulations of hybridizations that produced the tetraploid *C. davyi.* Each panel A-D represents a different pair of hybridizing lineages (species) listed above each graph. Within each panel, each curve corresponds to different assumed sub-genome dominance values (“Dom”)—see key in panel A. X-axis in each part corresponds to different possible hybridization times before present, expressed as RTU.

### *C. tenella*

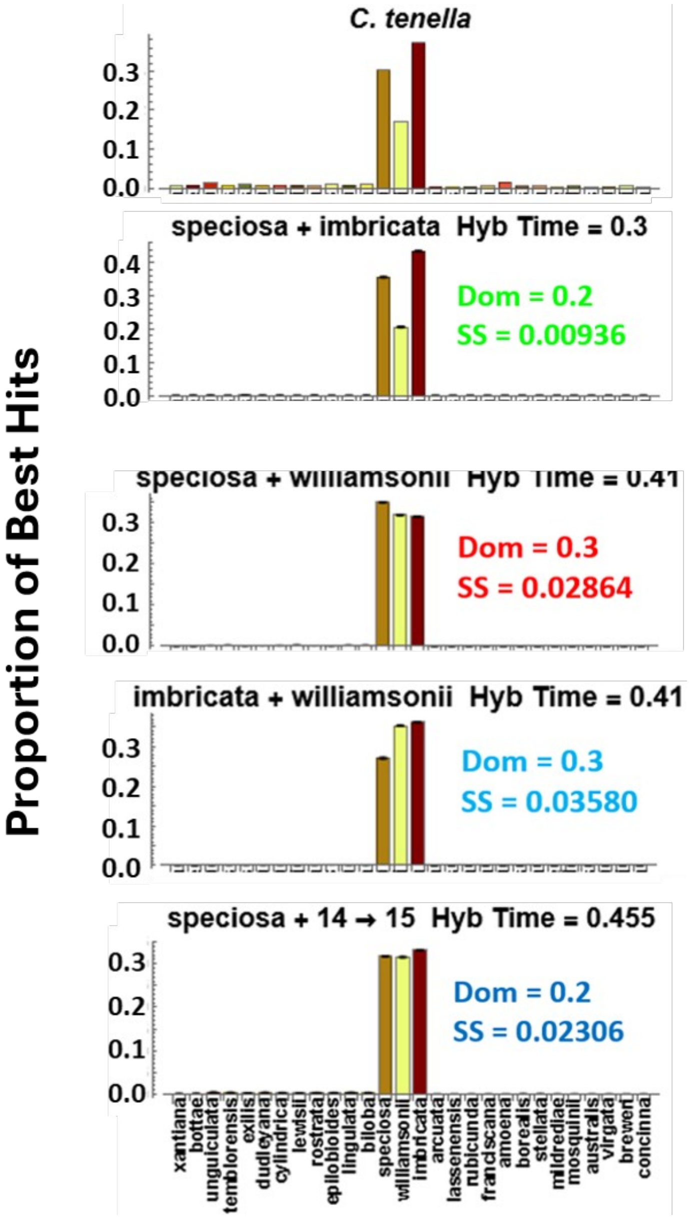


**Figure SIII.5**. Observed PBH spectrum for *C. tenella,* compared to simulated PBH spectra for each pair of possible progenitors examined. “Dom” is the value of subgenome dominance for the model. SS is Sum of Squared deviations for the model.

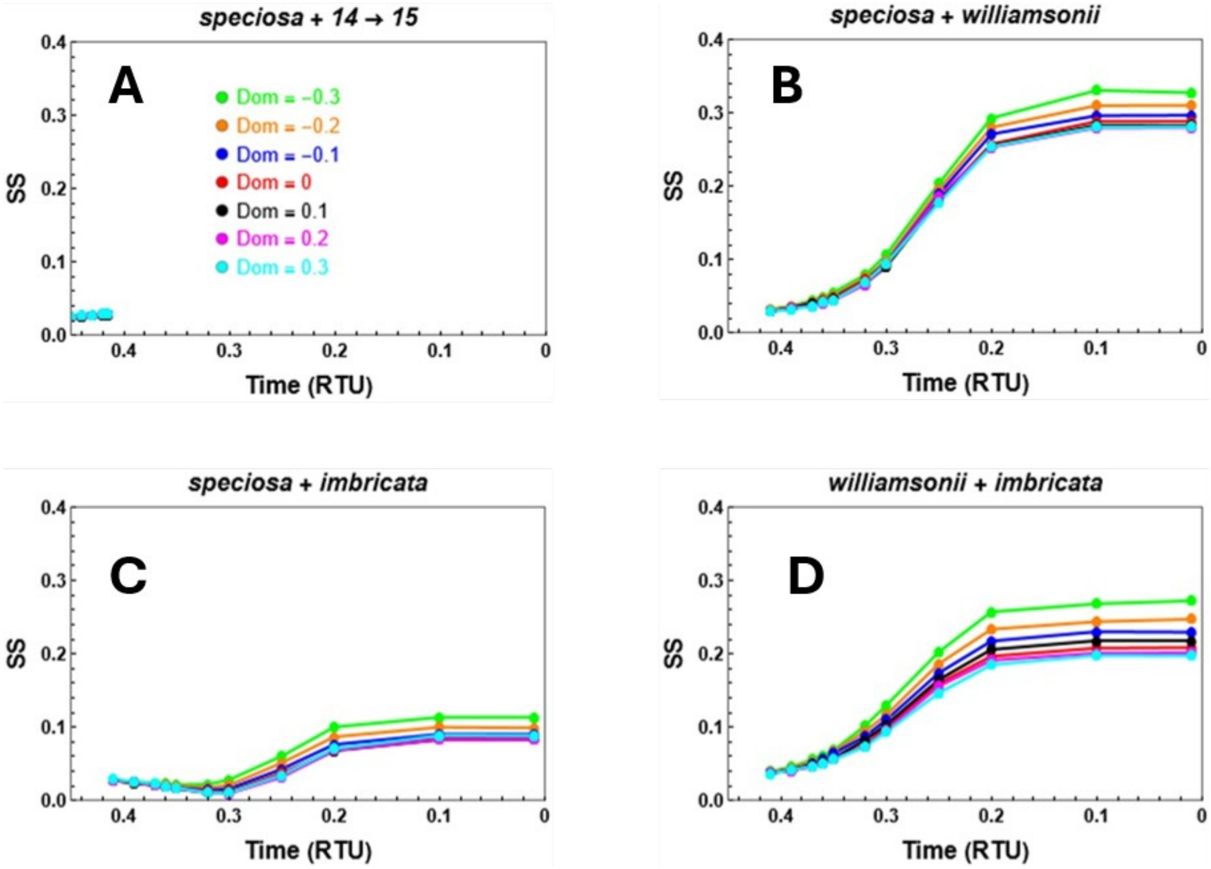


**Figure SIII.6**. Sums of Squared deviations (SS) between observed PBH spectra and predicted PBH for simulations of hybridizations that produced the tetraploid *C. davyi.* Each panel A-D represents a different pair of hybridizing lineages (species) listed above each graph. Within each panel, each curve corresponds to different assumed sub-genome dominance values ("Dom")—see key in panel A. X-axis in each part corresponds to different possible hybridization times before present, expressed as RTU.

### *C. similis*

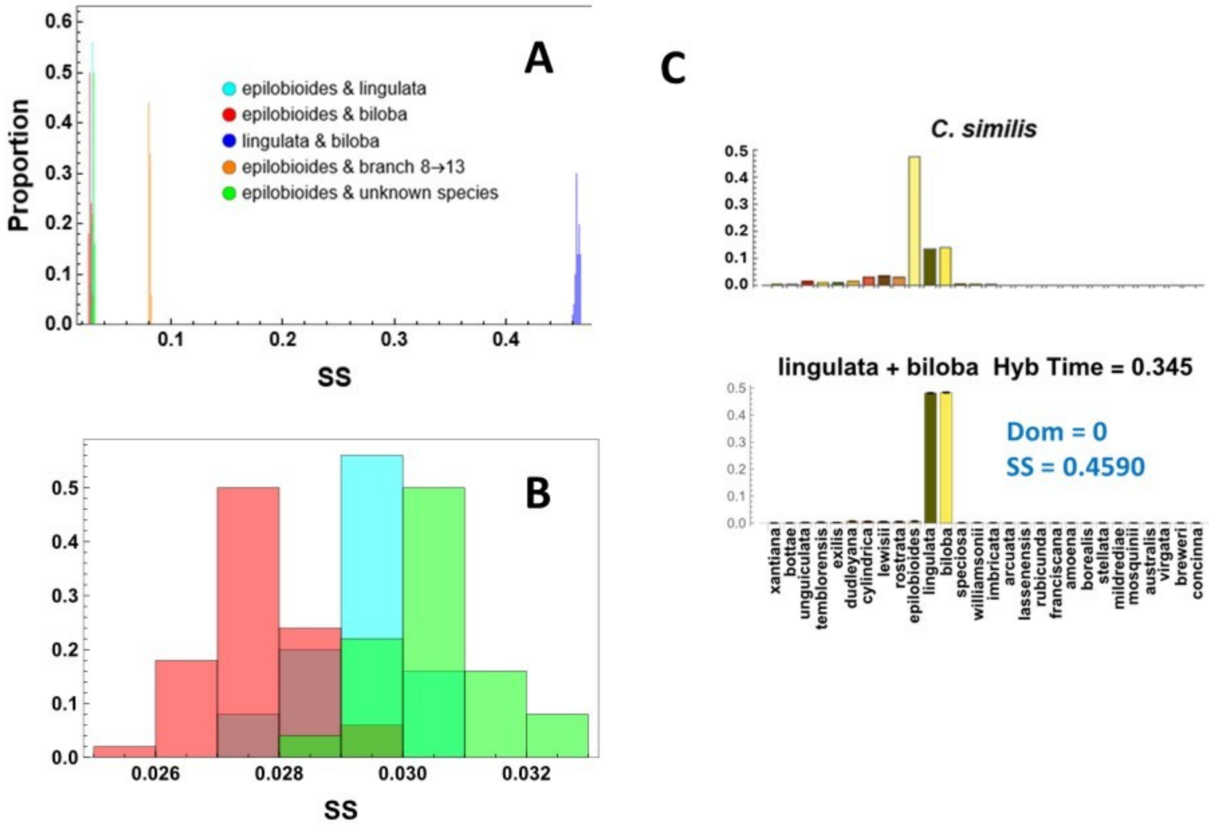


**Figure SIII.7.** A. Proportion of replicate simulations yielding binned Sum of Squared Deviations (SS) values for best-fitting model for each pair of possible progenitors of *C. similis*. B. Zoomed view of A. for low SS values, showing overlapping SS distributions for three different progenitor pairs. C. PBH spectrum for best-fitting model for progenitors *C. lingulata* and *C. biloba,* showing poor fit to observed PBH spectrum. "Dom" is value of subgenome dominance for the model. SS is Sum of Squared deviations for the model.

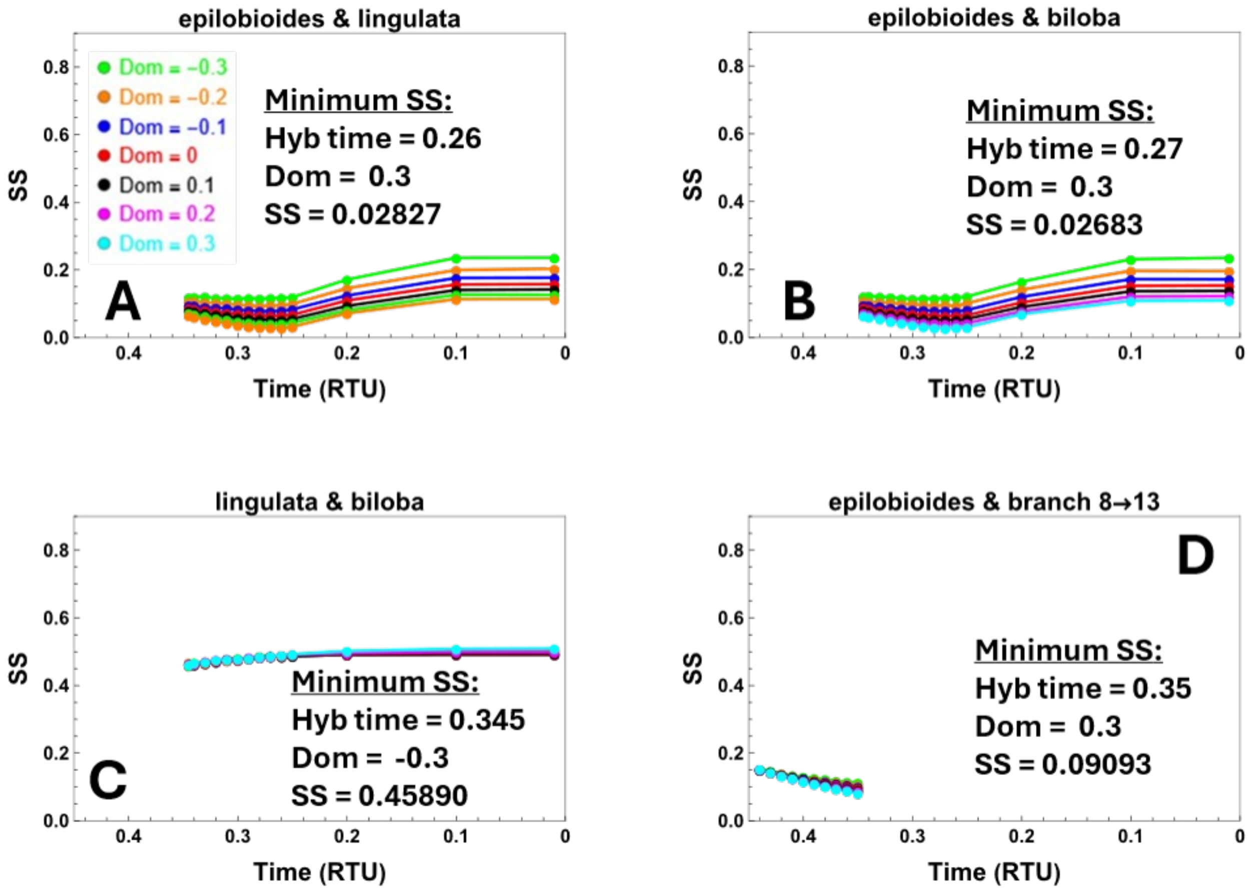


**Figure SIII.8.** Sums of Squared deviations (SS) between observed PBH and predicted PBH for models of hybridizations involving extant species that produced the tetraploid *C. similis.* Each part (A – D) represents a different pair of hybridizing lineages (species) listed above each graph. Within each panel, each curve corresponds to different assumed sub-genome dominance values ("Dom")—see key in panel A. X-axis in each panel corresponds to different possible hybridization times before present, expressed as RTU.

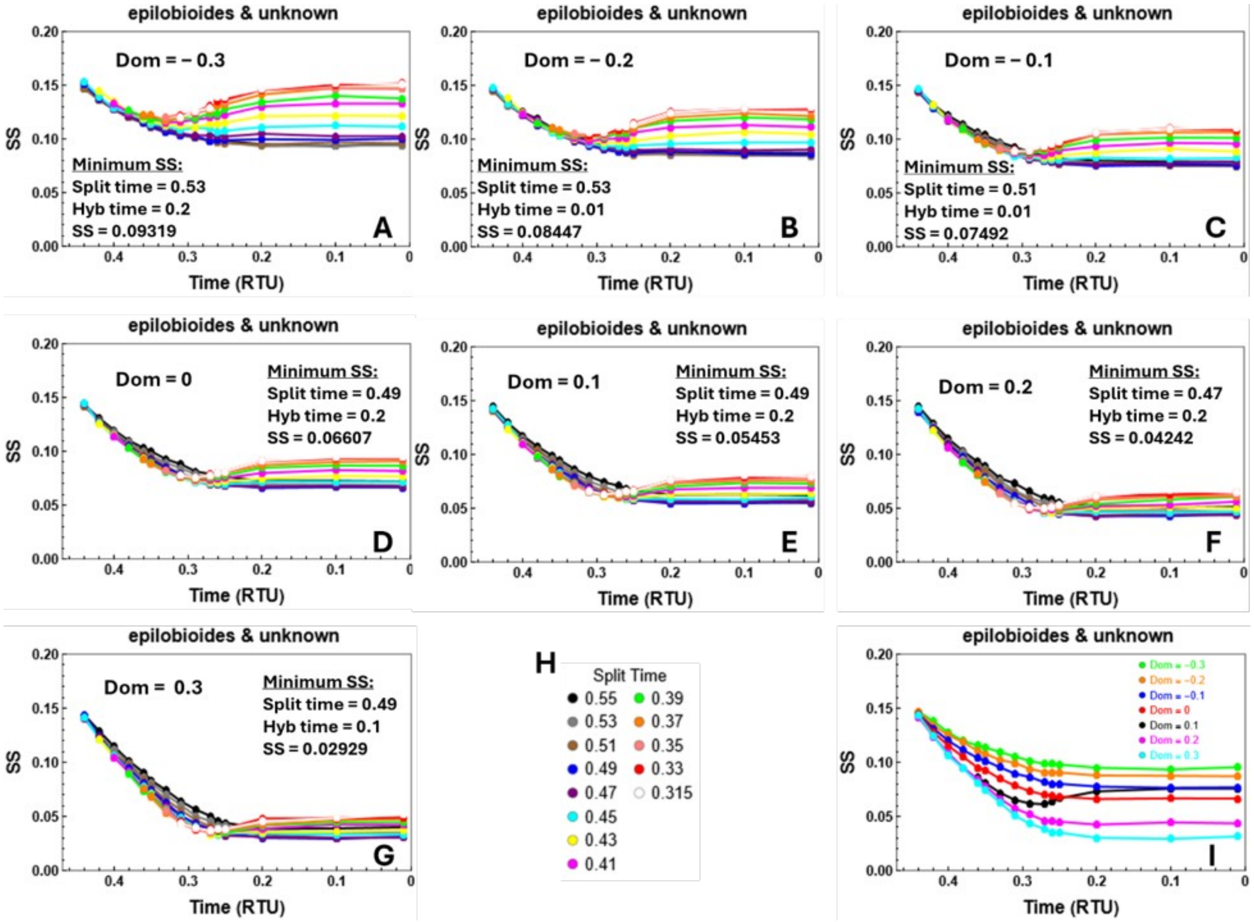


**Figure SIII.9.** Sums of Squared deviations (SS) between observed PBH and predicted PBH for model of hybridization between *C. epilobioides* and an unknown species (Fig. 3A) that produced the tetraploid *C. similis.* Panels A – G display model fit (SS) at different levels of subgenome dominance, where replicated lines show different split times between *C. epilobioides* and unknown species (see key in H). X-axis in each part corresponds to different possible hybridization times before present, expressed as RTU. **H.** Split time key for panels A-G. **I.** Sums of Squared deviations for the curves in panels A-G with the lowest SS under different levels of subgenome dominance listed in the key.

### *C. rhomboidea*

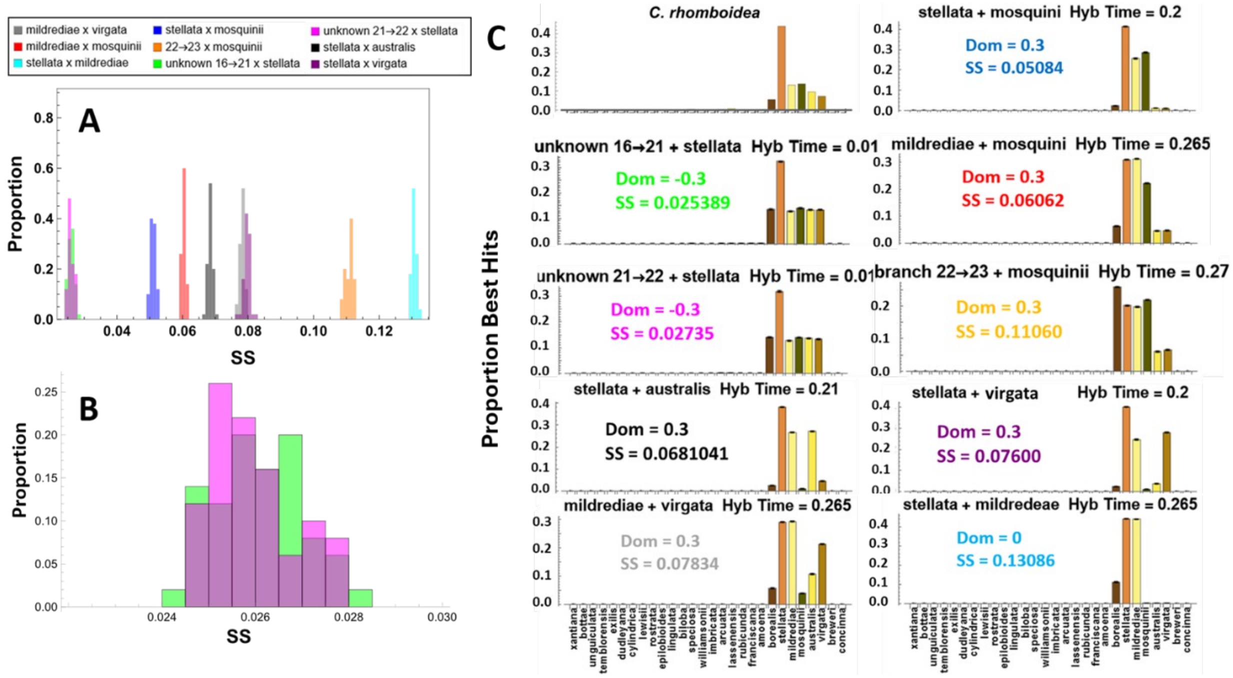


**Figure SIII.10.** A. Proportion of replicate simulations yielding binned Sum of Squared Deviations (SS) values for best-fitting model for each pair of possible progenitors for *C. rhomboidea*. **B.** Zoomed view of panel A. for low SS values. **C.** Observed PBH spectrum for *C. rhomboidea* and simulated PBH spectra for best-fitting model for each pair of possible progenitors examined. “Dom” is value of subgenome dominance for the model. SS is Sum of Squared deviations for the model.

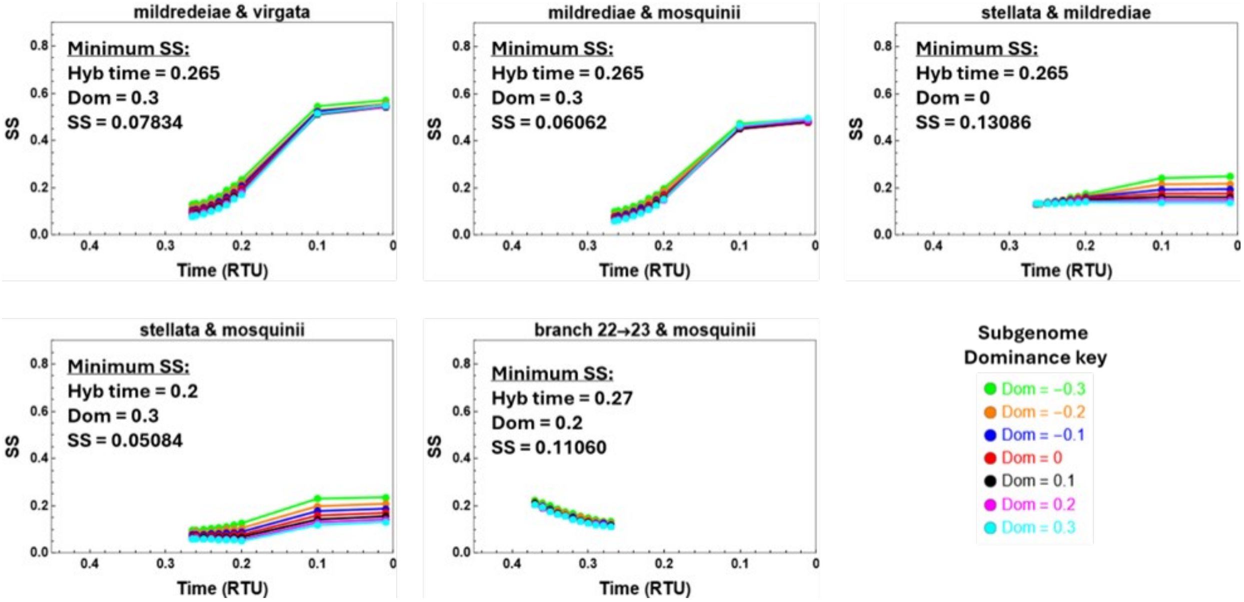


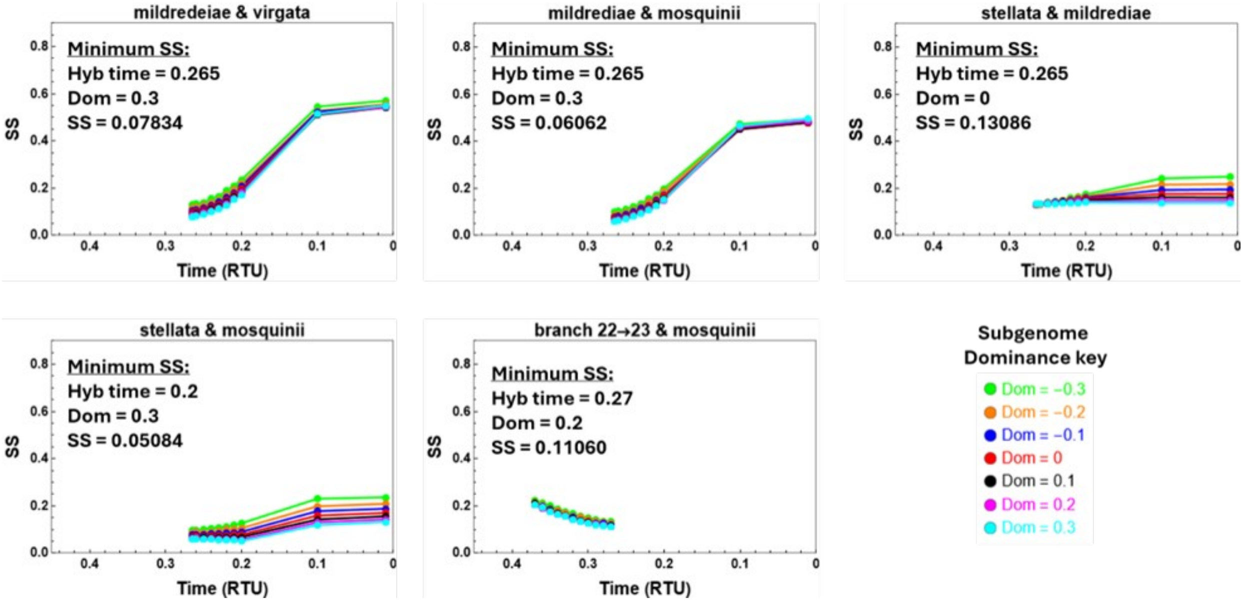


**Figure SIII.11.** Sums of Squared deviations (SS) between observed PBH and predicted PBH for models of hybridizations involving extant species that produced the tetraploid *C. rhomboidea.* Each panel represents a different pair of hybridizing lineages (species) listed above each graph. Within each panel, each curve corresponds to different assumed subgenome dominance values ("Dom")—see key in lower right. X-axis in each part corresponds to different possible hybridization times before present, expressed as RTU.

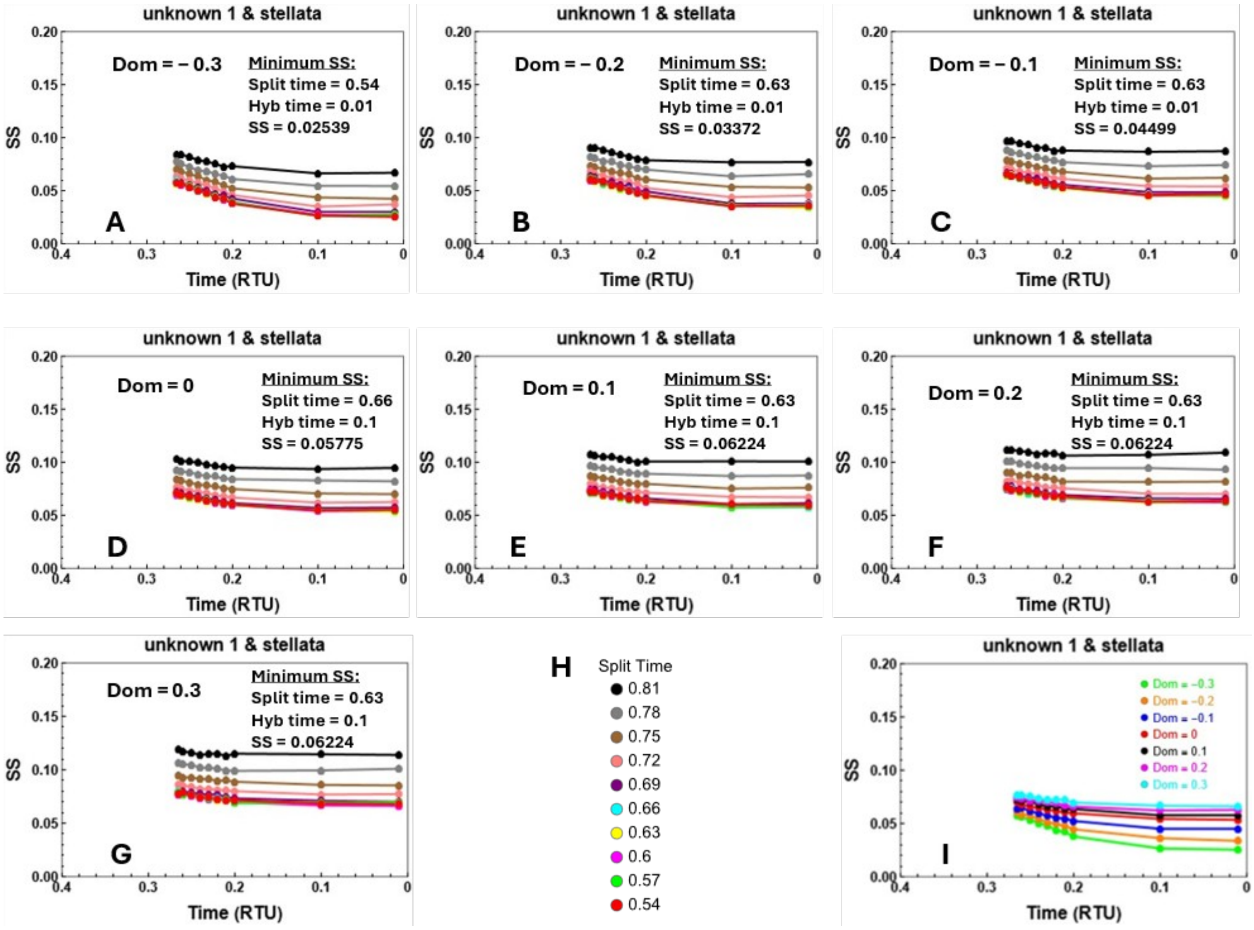


**Figure SIII.12.** Sums of Squared deviations (SS) between observed PBH and predicted PBH for simulations of hybridization between *C. stellata* and unknown species 1 (Fig. 3A) that produced the tetraploid *C. rhomboidea.* Panels A – G display model fit (SS) with different levels of subgenome dominance, where replicated lines show different split times between branch 16→21 and unknown species 1 (see key in H). X-axis in each part corresponds to different possible hybridization times before present, expressed as RTU. **H.** Split time key for panels A-G. **I.** Sums of Squared deviations for the curves in panels A-G with the lowest SS under different levels of subgenome dominance listed in the key.

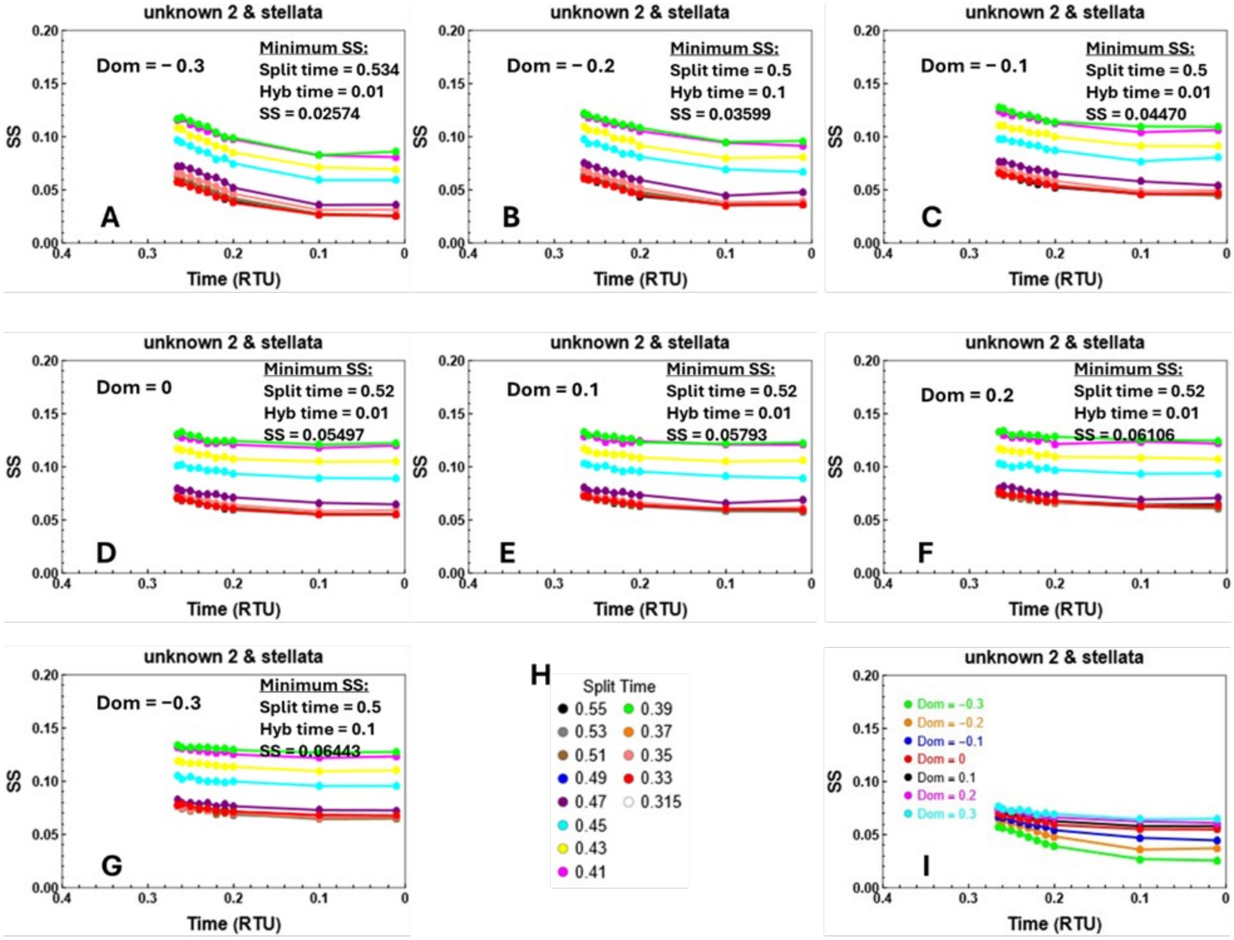


**Figure SIII.13.** Sums of Squared deviations (SS) between observed PBH and predicted PBH for simulations of hybridization between *C. stellata* and unknown species 2 (Fig. 3A) that produced the tetraploid *C. rhomboidea.* Panels A – G display model fit (SS) with different levels of subgenome dominance, where replicated lines represent different split times between branch 21→22 and unknown species 2 (see key in H). X-axis in each part corresponds to different possible hybridization times before present, expressed as RTU. **H.** Split time key for panels A-G. **I.** Sums of Squared deviations for the curves in panels A-G with the lowest SS under different levels of subgenome dominance listed in the key.

### *C. pulchella*

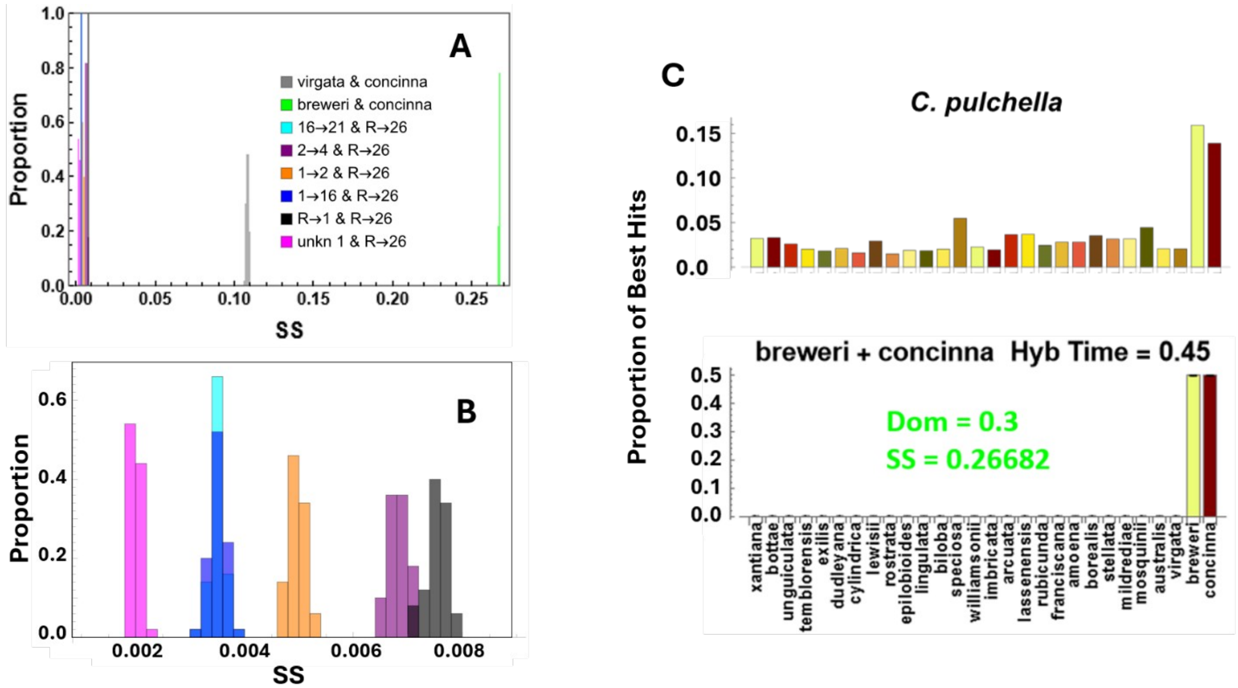


**Figure SIII.14.** A. Proportion of replicate simulations yielding binned Sum of Squared Deviations (SS) values for the best-fitting model for each pair of possible progenitors for *C. pulchella*. **B.** Zoomed view of panel A. for low SS values. **C.** Observed PBH spectrum for *C. pulchella* and simulated PBH spectra for the best-fitting model for progenitors *C. breweri* and *C. concinna,* showing poor fit to observed PBH spectrum. “Dom” is value of subgenome dominance for the model. SS is Sum of Squared deviations for the model.

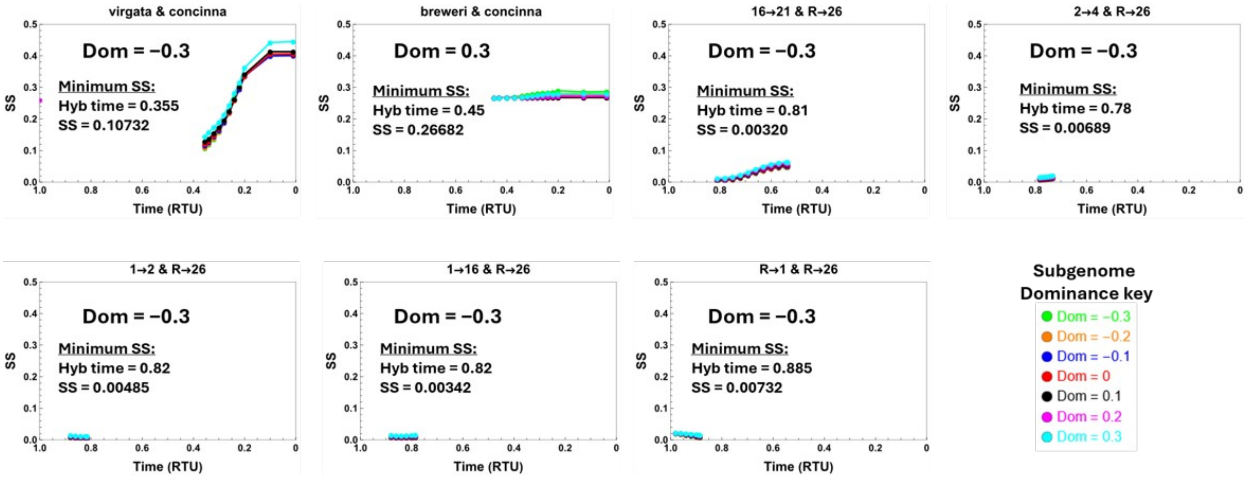


**Figure SIII.15.** Sums of Squared deviations (SS) between observed PBH and predicted PBH for simulations of hybridizations involving extant species that produced the tetraploid *C. pulchella.* Each panel represents a different pair of hybridizing lineages (species) listed above each graph. Within each panel, each curve corresponds to different assumed subgenome dominance values ("Dom")—see key in lower right. X-axis in each part corresponds to different possible hybridization times before present, expressed as RTU.

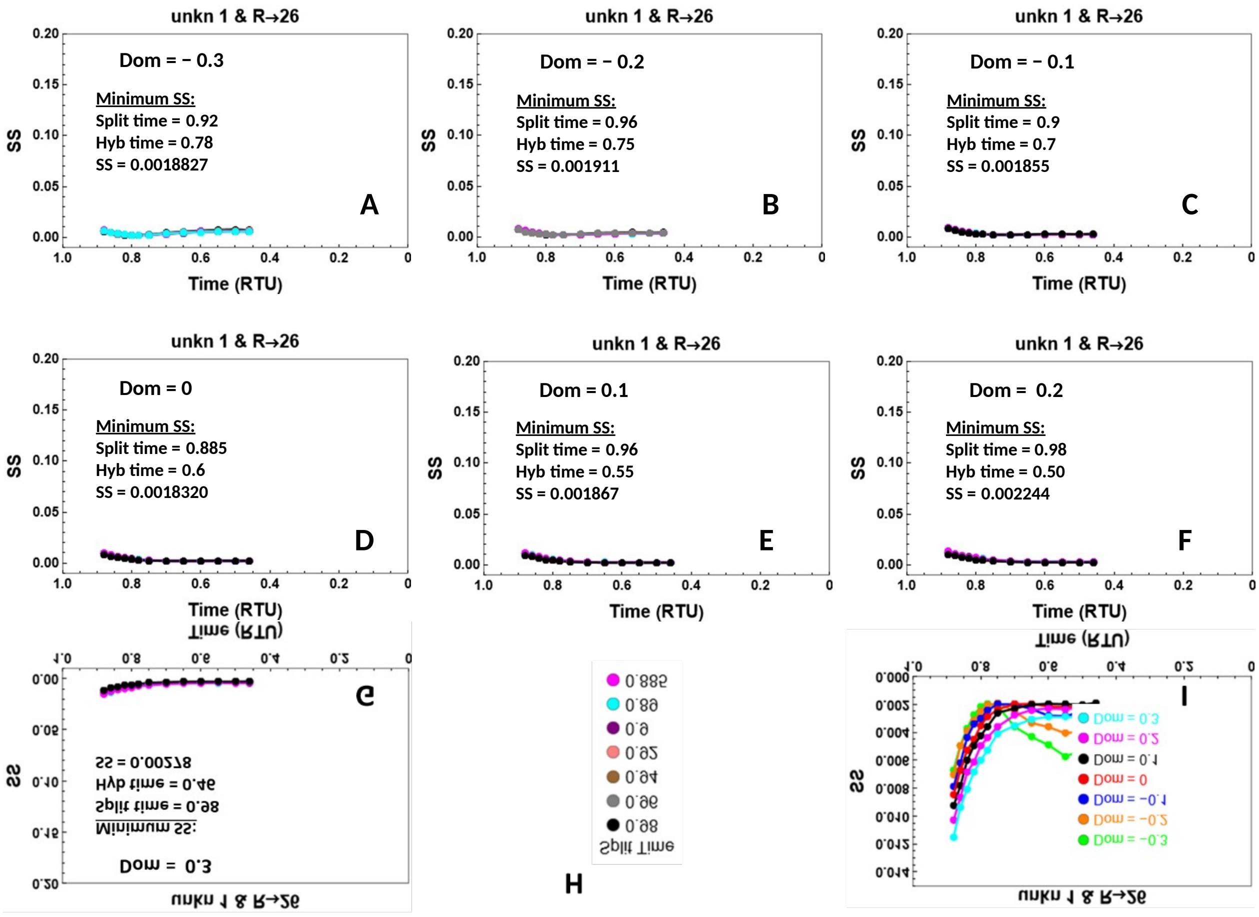


**Figure SIII.16.** Sums of Squared deviations (SS) between observed PBH and predicted PBH for simulations of hybridization between branch R→26 and an unknown species (Fig. 4A) that produced the tetraploid *C. pulchella.* Panels A – G display model fit (SS) with different levels of subgenome dominance, where replicated lines represent different split times between branch R→1 and an unknown species (see key in H). X-axis in each part corresponds to different possible hybridization times before present, expressed as RTU. **H.** Split time key for panels A-G. **I.** Sums of Squared deviations for the curves in panels A-G with the lowest SS under different levels of subgenome dominance listed in the key.

### IV. Supplementary Tables

**Table SIV.1.** Species name, sample ID, herbarium collection number, and seed source/original collection location for all samples used in this study.

| Species | Sample | Collection Number | Source | Collection Location |
|---|---|---|---|---|
| Clarkia amoena ssp. amoena | AmoeAmoe_MR8 | | Larner Seeds Nursery | |
| Clarkia arcuata | Arcu_8812 | RSA495974 | Weeden Lab, Montana State University | Mariposa County, CA |
| Clarkia australis | Aust_2 | | Moeller Lab, University of Minnesota | Tuolumne County, CA |
| Clarkia biloba ssp. biloba | BiloBilo_15880 | RSA495266 | California Botanical Garden | Tuolumne County, CA |
| Clarkia borealis | Boreal_4911 | | Weeden Lab, Montana State University | Shasta County, CA |
| Clarkia bottae | Bott_5-1-2 | | Wild Collected | Santa Barbara County, CA |
| Clarkia breweri | Brew_15883 | RSA495954 | California Botanical Garden | Stanislaus County, CA |
| Clarkia concinna ssp. concinna | ConcConc_45266 | | US Department of Agriculture GRIN | |
| Clarkia cylindrica ssp. cylindrica | CylCyl_1-3-1 | | Wild Collected | Los Angeles County, CA |
| Clarkia davyi | Davy_8417 | | Weeden Lab, Montana State University | Mendocino County, CA |
| Clarkia delicata | Deli_9605 | DAV158368 | Weeden Lab, Montana State University | San Diego County, CA |
| Clarkia delicata | Deli_9611 | DAV158371 | Weeden Lab, Montana State University | San Diego County, CA |
| Clarkia dudleyana | Dudl_24443 | | California Botanical Garden | Los Angeles County, CA |
| Clarkia epilobioides | Epilo_21078 | RSA678282 | California Botanical Garden | Riverside County, CA |
| Clarkia exilis | Exil_46 | | Mazer Lab, UC Santa Barbara | Kern County, CA |
| Clarkia franciscana | Franc_OAK | | UC Berkeley Botanical Garden | Alameda County, CA |
| Clarkia imbricata | Imbric_15893 | | California Botanical Garden | Sonoma County, CA |
| Clarkia lassenensis | Lass_RCL2 | | Weeden Lab, Montana State University | Modoc County, CA |
| Clarkia lewisii | Lewi_45494 | | US Department of Agriculture GRIN | |
| Clarkia lingulata | Ling_21442 | | California Botanical Garden | Merced County, CA |
| Clarkia mildrediae | Mild_8759 | DAV138173 | Weeden Lab, Montana State University | Butte County, CA |
| Clarkia mosquinii | Mosq_9312 | | Weeden Lab, Montana State University | Butte County, CA |
| Clarkia pulchella | Pulch_37388 | | US Department of Agriculture GRIN | Idaho |
| Clarkia pulchella | Pulch_39037 | | US Department of Agriculture GRIN | |
| Clarkia rhomboidea | Rhomb_23-1-1 | | Wild Collected | Kern County, CA |
| Clarkia rhomboidea | Rhomb_7503 | | Kay Lab, UC Santa Cruz | Kern County, CA |
| Clarkia rhomboidea | Rhomb_WW2 | | Wilamette Wildings Nursery | Oregon |
| Clarkia rostrata | Rost_15899 | RSA495300 | California Botanical Garden | Mariposa County, CA |
| Clarkia rubicunda | Rubi_20733 | | California Botanical Garden | Santa Clara County, CA |
| Clarkia similis | Sim_9610 | DAV158473 | Weeden Lab, Montana State University | San Diego County, CA |
| Clarkia speciosa ssp. speciosa | SpecSpec_15903 | RSA495269 | California Botanical Garden | San Luis Obispo County, CA |
| Clarkia stellata | Stell_9317 | | Weeden Lab, Montana State University | Plumas County, CA |
| Clarkia tembloriensis ssp. calientensis | TemblCal_1-1 | | UC Berkeley Botanical Garden | Kern County, CA |
| Clarkia tenella | Tene_MR5 | | Seedhunt Nursery | |
| Clarkia unguiculata | Unguic_45269 | | US Department of Agriculture GRIN | |
| Clarkia virgata | Virg_2 | | Moeller Lab, University of Minnesota | Tuolumne County, CA |
| Clarkia virgata | Virg_3 | | Moeller Lab, University of Minnesota | Tuolumne County, CA |
| Clarkia williamsonii | Willia_18946 | | California Botanical Garden | Madera County, CA |
| Clarkia xantiana ssp. xantiana | XantXant_MDR1 | | Eckhart Lab, Grinnell College | Kern County, CA |

**Table SIV.2**. Observed Proportion Best Hits (PBH) to each diploid species from each tetraploid species. Values are averaged between replicates (if available).

| | Observed Proportion Best Hits (PBH) | | | | | |
|---|---|---|---|---|---|---|
| | Polyploid Species | | | | | |
| *Diploid Species* | *C. davyi* | *C. tenella* | *C. delicata* | *C. similis* | *C. rhomboidea* | *C. pulchella* |
| *C. xantiana* | 0.0053006 | 0.005052546 | 0.002531564 | 0.007974259 | 0.003122832 | 0.032279482 |
| *C. bottae* | 0.0059716 | 0.006871463 | 0.003095368 | 0.007204813 | 0.004005132 | 0.033040993 |
| *C. unguiculata* | 0.0119431 | 0.01401240 | 0.037714652 | 0.019515949 | 0.003262861 | 0.025910563 |
| *C. temblorensis* | 0.0074477 | 0.007477769 | 0.042894029 | 0.014199776 | 0.002357362 | 0.020168183 |
| *C. exilis* | 0.005770 | 0.008218809 | 0.402790779 | 0.014619474 | 0.002684782 | 0.018141636 |
| *C. dudleyana* | 0.0073135 | 0.00767987 | 0.005328512 | 0.019236150 | 0.003057674 | 0.020824761 |
| *C. cylindrica* | 0.006240 | 0.00767987 | 0.014152113 | 0.034135422 | 0.002133587 | 0.015985749 |
| *C. lewisii* | 0.008387 | 0.00794934 | 0.01708559 | 0.037493005 | 0.002108713 | 0.029184488 |
| *C. rostrata* | 0.0043612 | 0.005726219 | 0.014917422 | 0.033715725 | 0.002046163 | 0.014832530 |
| *C. epilobioides* | 0.0066425 | 0.008353544 | 0.43095723 | 0.475447678 | 0.002503806 | 0.018714491 |
| *C. lingulata* | 0.0073806 | 0.008084074 | 0.005162236 | 0.136751539 | 0.003162331 | 0.018372781 |
| *C. biloba* | 0.0064412 | 0.008218809 | 0.00539294 | 0.141927812 | 0.002796324 | 0.020168809 |
| *C. speciosa* | 0.2861648 | 0.302344382 | 0.00163137 | 0.009443201 | 0.004285477 | 0.054741872 |
| *C. williamsonii* | 0.1514359 | 0.169765562 | 0.002264535 | 0.007974259 | 0.002414622 | 0.022592696 |
| *C. imbricata* | 0.4276033 | 0.373012665 | 0.001466738 | 0.005106323 | 0.002339216 | 0.019358552 |
| *C. arcuata* | 0.0019458 | 0.002627324 | 0.000800537 | 0.003637381 | 0.004013243 | 0.036489452 |
| *C. lassenensis* | 0.0030864 | 0.00357047 | 0.000765309 | 0.002308338 | 0.007096904 | 0.036894859 |
| *C. rubicunda* | 0.0013419 | 0.002425222 | 0.000598485 | 0.002028539 | 0.003078963 | 0.024398738 |
| *C. franciscana* | 0.0038916 | 0.00511991 | 0.000965169 | 0.004196978 | 0.002996672 | 0.028120761 |
| *C. amoena* | 0.0110709 | 0.015022905 | 0.000665653 | 0.002168439 | 0.003549471 | 0.028124515 |
| *C. borealis* | 0.0035561 | 0.004985179 | 0.001129801 | 0.003287633 | 0.055149571 | 0.035166909 |
| *C. stellata* | 0.003489 | 0.004378874 | 0.001064277 | 0.003707331 | 0.439907931 | 0.031512338 |
| *C. mildredeae* | 0.0029522 | 0.004176772 | 0.001096765 | 0.002867935 | 0.131163106 | 0.031589178 |
| *C. mosquinii* | 0.0036232 | 0.004985179 | 0.001265234 | 0.003987129 | 0.13709505 | 0.044429513 |
| *C. australis* | 0.0008052 | 0.001549448 | 0.000466341 | 0.001329043 | 0.095279048 | 0.020475541 |
| *C. virgata* | 0.0027509 | 0.002896793 | 0.000565997 | 0.001468942 | 0.072248836 | 0.020441815 |
| *C. breweri* | 0.0112721 | 0.005860954 | 0.002531564 | 0.003007834 | 0.003256953 | 0.159148411 |
| *C. concinna* | 0.0018116 | 0.001953651 | 0.000699785 | 0.001259093 | 0.00288337 | 0.138890382 |
